# Gαi2 Interaction with EB1 Controls Microtubule Dynamics and Rac1 Activity in *Xenopus* Neural Crest Cell Migration

**DOI:** 10.1101/2023.09.07.556733

**Authors:** Soraya Villaseca, Juan Ignacio Leal, Lina Mariana Tovar, María José Ruiz, Jossef Guajardo, Hernan Morales-Navarrete, Roberto Mayor, Marcela Torrejón

**Affiliations:** Laboratory of Signaling and Development (LSD), Group for the Study of Developmental Processes (GDeP), Department of Biochemistry and Molecular Biology, Faculty of Biological Sciences, Universidad de Concepción, Casilla 160-C, Concepción, Chile; Department of Systems Biology of Development, University of Konstanz, Germany; Department of Cell and Developmental Biology, University College London, WC1E 6BT London, England, UK

**Author notes:** Correspondence to Marcela Torrejón or Roberto Mayor.

**Keywords:** Gαi2, microtubules, cell migration, heterotrimeric G-Protein, focal adhesion, cell polarity, protrusion formation, neural crest

## Abstract

Cell migration is a complex and essential process in various biological contexts, from embryonic development to tissue repair and cancer metastasis. Central to this process are the actin and tubulin cytoskeletons, which control cell morphology, polarity, focal adhesion dynamics, and overall motility in response to diverse chemical and mechanical cues. Despite the well-established involvement of heterotrimeric G proteins in cell migration, the precise underlying mechanism remains elusive, particularly in the context of development. This study explores the involvement of Gαi2, a subunit of heterotrimeric G proteins, in cranial neural crest cell migration, a critical event in embryonic development. Our research uncovers the intricate mechanisms underlying Gαi2 influence, revealing a direct interaction with the microtubule-associated protein EB1, and through this with tubulin, suggesting a regulatory function in microtubule dynamics modulation. Here, we show that Gαi2 knockdown leads to microtubule stabilization, alterations in cell polarity and morphology with an increased Rac1-GTP concentration at the leading edge and cell-cell contacts, impaired cortical actin localization and focal adhesion disassembly. Interestingly, in Gαi2 knockdown cells, RhoA-GTP was found to be reduced at cell-cell contacts and concentrated at the leading edge, providing evidence of Gαi2 significant role in polarity. Remarkably, treatment with nocodazole, a microtubule-depolymerizing agent, effectively reduces Rac1 activity, restoring cranial NC cell morphology, actin distribution, and overall migration. Collectively, our findings shed light on the intricate molecular mechanisms underlying cranial neural crest cell migration and highlight the pivotal role of Gαi2 in orchestrating microtubule dynamics through EB1 and EB3 interaction, modulating Rac1 activity during this crucial developmental process.

## Introduction

Cell migration plays a crucial role throughout an organism’s lifespan. During embryogenesis, it drives various processes such as germ layer formation and organogenesis. In adults, proper wound healing, immune response and tissue homeostasis rely on accurate cell migration, while abnormal cell migration is associated with severe pathologies, including cancer metastasis (Chang et al., 2013; Mayor and Etienne-Manneville, 2016; Ridley et al., 2003; Vedula et al., 2013; Trainor P.A., 2010). The process of cell migration is orchestrated by a cascade of molecular and morphological events. These events encompass cell polarization, protrusion generation, substrate adhesion, cell body translocation through contraction and release of adhesions, ultimately leading to cell detachment (Ridley et al., 2003; Abercrombie et al., 1971; Heath and Dunn, 1978; Huttenlocher et al., 1995; Lauffenburger and Horwitz, 1996). Actin microfilaments and microtubules are crucial for cell shape, cell adhesion and motility, and their remodeling is vital during these events (Akhshi et al., 2013; Garcin and Straube, 2019). The crosstalk between actin microfilaments and microtubules involves small GTPases from the Rho family and other components of the cytoskeletal networks. These factors govern cell polarity, actin polymerization and actomyosin contractility (Seetharaman and Etienne-Manneville, 2020; Garcin and Straube, 2019; Bouchet and Akhmanova, 2017; Schaks et al., 2019). Additionally, microtubules participate in trafficking and deliver molecules to the leading edge of the migrating cell, influencing mechanosensing and regulating focal adhesion (FA) dynamics, migration velocity, persistence and directionality (Etienne-Manneville, 2013; Gu et al., 2011).

The neural crest (NC) has long served as a valuable model for studying cell migration during development and has contributed to our understanding of the chemical and mechanical cues that govern this process (Barriga et al., 2018; Kelleher et al., 2006; Mayor and Theveneau, 2013; Toro-Tapia et al., 2018). The NC, originating at the neural plate border, represents a transient embryonic cell population that undergoes extensive migration to differentiate into various tissues (Szabó et al., 2018). During the pre-migratory stage, cranial NC cells establish an epithelium with distinct apico-basal polarity. Following specification, these cells undergo an epithelial-mesenchymal transition (EMT), enabling them to acquire motility, lose their epithelial polarity and undergo a switch from the highly adhesive E-cadherin to the less adhesive N-cadherin. This transition facilitates their migration and colonization of adjacent tissues (Bronner-Fraser and Sauka-Spengler, 2008; Kuriyama and Mayor, 2008; Steventon and Mayor, 2012; Theveneau and Mayor, 2012). In polarized cranial NC cells, Rac1 and Cdc42 exhibit activity at the leading edge, facilitating the formation of cell protrusions (Matthews et al., 2008; Ridley et al., 2011; Liu et al., 2013). Conversely, RhoA activity at the rear of the cell regulates cell contraction (Theveneau et al., 2010; Shoval and Kalcheim, 2012). Furthermore, in *Xenopus*, Par3 plays a negative regulatory role in Rac1 activity at cell-cell contacts by inhibiting Rac-GEF Trio, thereby inducing microtubules catastrophe during cranial NC cell migration (Moore et al., 2013). As previously mentioned, the interaction between actin and microtubules is mediated by small GTPases from the Rho family (Rodriguez et al., 2003).

Numerous studies have demonstrated that heterotrimeric G-protein-mediated signals are involved in cell migration, including in NC cells (Cotton and Claing, 2009; Nobes and Hall, 1995; Kjoller and Hall, 1999; Sah et al., 2000; Rohde and Heisenberg, 2007; Fuentealba et al., 2013; Toro-Tapia et al., 2018; Leal et al., 2018). Recent studies have demonstrated that Ric-8A, a Guanine Nucleotide Exchange Factor (GEF) for Gα subunits of heterotrimeric G proteins, plays a regulatory role in NC cell migration by influencing polarity properties (Leal et al., 2018) and facilitating focal adhesion formation through Gα13 subunit (Fuentealba et al., 2013; Toro-Tapia et al., 2018). Nevertheless, all the Gα family subunits from heterotrimeric G proteins have been implicated in cell migration (Cotton and Claing, 2009), and their expression has been observed in the NC region of *Xenopus* embryos (Fuentealba et al., 2016). Specifically, the Gαi/o family has been strongly associated with cell migration through chemotactic signaling in other tissues (Han et al., 2005; Hwang et al., 2007; Pero et al., 2007; Zarbock et al., 2007). Treatment with pertussis toxin (PTX), an inhibitor of Gαi/o family protein function, results in impaired chemotaxis in macrophages (Wiege et al., 2012). Gαi2 is expressed in NC cells (Fuentealba et al., 2016) and it has been implicated in the regulation of cell migration in other tissues through a GPCR-dependent mechanism (Hwang et al., 2007; Hwang et al., 2015). Additionally, studies conducted on Gαi2 knockout mice demonstrate a significant reduction in the macrophages and neutrophils recruitment to inflammatory sites (Han et al., 2005; Zarbock et al., 2007; Wiege et al., 2012). Notably, Gαi subunit has been reported to interact with microtubules motor components such as dynein and polarity molecules, thereby influencing asymmetric cell division controlling the spindle positioning in a GPCR independent mechanism (Kiyomitsu T., 2019; Schaefer et al., 2001; Yu et al., 2000; Villaseca et al., 2022). Despite the established role of Gαi2 in chemotaxis through GPCR activation, limited information is available regarding its specific mechanism by which it controls cell migration (Villaseca et al., 2022). Hence the primary aim of this study was to unravel the mechanism by which Gαi2 controls cranial NC cell migration using *Xenopus* as an *ex vivo* model.

Our findings highlight the essential role of Gαi2 in microtubules dynamic to modulate the establishment of accurate polarity proteins and focal adhesion disassembly, thereby facilitating NC cell migration. Under Gαi2 knockdown conditions, we observed a decreased in microtubules dynamics and an increase in Rac1 activity. In this sense, our findings showed a physical and functional interplay between Gαi2 with microtubule accessory proteins, such as EB1 and EB3, regulating microtubules dynamic and their components, possible controlling downstream effector like Rac1. Therefore, regulating actin cytoskeleton, protrusion formation and focal adhesion disassembly. Consistently, we showed that low concentrations of nocodazole reduce Rac1 activity in Gαi2 depleted cells and restore cranial NC cell morphology, actin distribution and consequently cell migration. This study offers new insights into the previously unrecognized function of Gαi2 in regulating microtubule dynamics, through its interaction with EB1/EB3 to ensure proper polarization, cell morphology and focal adhesion dynamics during cell migration.

## Results

### Gαi2 is required for cranial neural crest cell migration

Since cranial NC cells express Gαi2 during migration stages (Fuentealba et al., 2016), we analyzed whether it plays any role on NC migration. To achieve this, we performed loss-of-function studies by injecting a translational inhibitory antisense morpholino oligonucleotide against to Gαi2 (Gαi2MO), in *Xenopus* embryos, followed by WISH against the NC marker *snail2*. The injection of Gαi2MO or controlMO into one dorsal and one ventral blastomere of eight-cells stage embryo did not interfere with NC induction, as the distribution of *snail2* expressing cells remained unaffected at stage 16 (**Fig. 1A**). To evaluate cranial NC cell migration, we performed WISH against *snail2* and quantified the phenotype by measuring the total length of the three migratory routes (mandibular, hyoid and branchial) at stage 22-23, as depicted in (**Fig. 1F)**. The results showed severe alterations in migration, with the localization of *snail2* positive cells affected at stage 22-23 (**Fig. 1B,C,E,G**) in 60-70% (n=49) of injected embryos from *X. tropicalis*, showing impaired NC cell migration. Importantly, we demonstrated that migration defects can be rescued in Gαi2MO embryos (n=31) upon co-injection with Gαi2 mRNA that does not bind the Gαi2MO, indicative that the morphant phenotype observed is specific (**Fig. 1D,G)**. Western blot analysis of Gαi2 shows that the Gαi2MO significantly reduced the levels of the protein (**Fig. 1H,I**). It is important to note that similar results were observed in *Xenopus laevis* (**supplementary material Fig. S1A-E**). In later developmental stages, such as stage 32, WISH revealed alterations in migration as well, albeit to a lesser extent compared to the early stages (22-23). This suggests a phenotype characterized by delayed migration (**supplementary material Fig. S1F-H**). In addition, the injection of Gαi2MO had no effect on Epithelial-Mesenchymal Transition (EMT), as recognized by the expression of E-cadherin and N-Cadherin, similar to control NC cells (**supplementary material Fig. S2A-F**). Together, these observations show that Gαi2 is required for a proper cranial NC cell migration *in vivo*.

**Figure 1.**
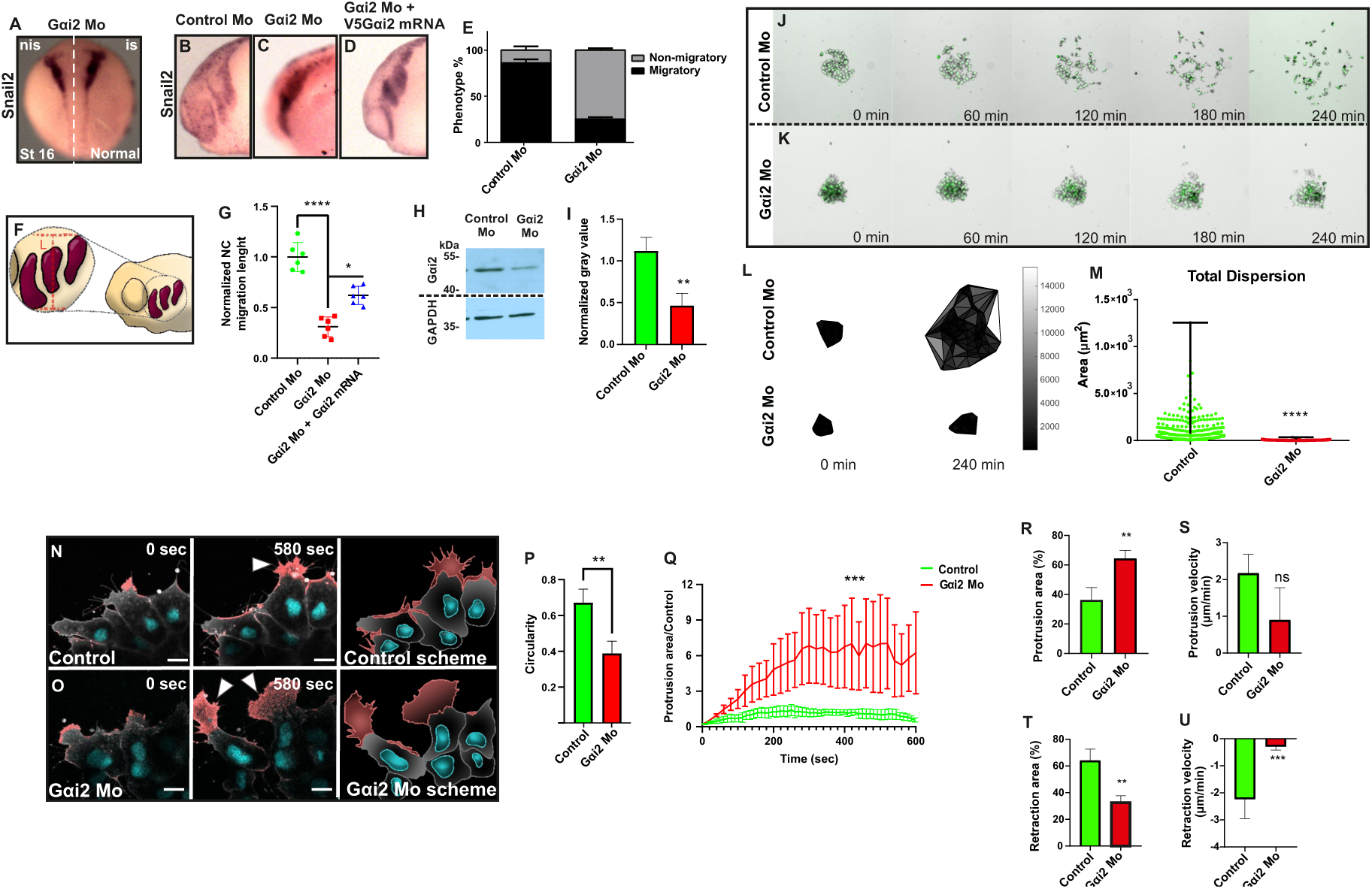
Gαi2 is required for NC migration *in vivo* and *in vitro*, and knockdown generates longer cells with bigger protrusions. **A-D.** *In situ* hybridization against *snail2* on *Xenopus tropicalis (X.t)* embryos at stage 16 to analyze *in vivo* NC induction n = 16 (A), and at stages 22-23 to study *in vivo* NC migration (B-D). n control = 49, n morphant = 49, n rescue = 31. nis: non-injected side, is: injected side. **E.** Percentage of migratory/non-migratory phenotype showing a dramatic impairment of NC cell migration in Gαi2 knockdown condition. **F.** Schematic representation of *Xenopus* embryos showing the cranial NC streams in purple, illustrating that the migration length was measured at the midpoint to the full width of the three cranial NC streams. **G.** Quantification of the migration length in B, C and D. Each point is the mean value of a total of 6 independent experiments (n= 3 embryos from each condition in each experiment). Error bars are ± s.e.m. **P≤0.01, ****P≤0.0001 (two-tailed Student’s t-test) **H-I.** Western blot of lysates from *Xenopus tropicalis* embryos injected with Gαi2 morpholino shows efficiently Gαi2 knockdown. Intensity was normalized to GAPDH to quantifying the decrease in Gαi2 expression induced by the morpholino. **J-K.** Explants of cranial NC cells were extracted from control embryos and embryos injected with Gαi2 morpholino, and migration was evaluated under the different conditions during 4 hours by time-lapse. **L-M.** Delaunay triangulation was performed to quantify cell dispersion. The average area from total dispersion after 4 hours was plotted. Error bar: s.e.m., ****P≤0.0001, n = 13 explants (Kruskal-Wallis non-parametric test). 10X magnification. **N-O.** Embryos were injected with membrane-GFP to analyze cell morphology. Scale bar: 10 μm. **P-Q.** Circularity and protruding area parameters were measured in 3 random cells for each experiment (n = 3 experiments per condition). Error bars correspond to ± s.e.m. **P≤0.01; ***P≤0.001 (T independent test). **R-U.** The percentage of protruding area, the percentage of retractable area, the protrusion growth rate and retraction rate were quantified using the ADAPT software from ImageJ. Error bars correspond to ± s.e.m. **P≤0.086; ***P≤0.001; ns: non-significant. T independent test.

To further characterize the effect of Gαi2MO, NC explants from stage 17 embryos were cultured on fibronectin, and their migratory behavior was then observed and recorded using time-lapse imaging. To quantify cell dispersion, we employed Delaunay triangulation (Carmona-Fontaine et al., 2011), revealing a severe impairment in Gαi2 morphant cells migration compared to control cells (**Fig. 1J-M**, **supplementary material movie S1**). As it is showed in (**Fig. 1J,K**), the injection of Gαi2MO resulted in a significant reduction in the area covered by cranial NC cells after 4 hours. Taken together these results support the notion that Gαi2 controls cranial NC cell migration.

### Gαi2 is required to regulate cell morphology during migration

To further explore the influence of Gαi2 during cranial NC cell migration, we conducted an analysis of the impact of Gαi2 knockdown on cell morphology. To achieve this, we used NC explants, which were injected with mRNAs encoding H2B-Cherry and membrane-GFP to label nuclei and cell membrane, respectively (**Fig. 1N,O**), and lifeactin-GFP to label actin cytoskeleton (**supplementary material Fig. S2G,H**). Cell morphology was evaluated under both control and morphant conditions, focusing on cell shape and protrusion dynamics (**Fig. 1N-Q, supplementary material Fig. S2G,H)**. We found that Gαi2 depleted NC cells exhibited larger and more stable protrusions when compared with control cells, accompanied by a notable reduction in cell circularity (**Fig. 1N-P**). The quantification of the protruding and retraction area showed that cranial NC cells with reduced Gαi2 exhibited a significant increase of approximately 70% in the protruding area whereas control cells only displayed a protruding area of 30% (**Fig. 1Q,R**). Conversely, the retractile area in the morphant cells decreased in comparison to the control cells, with values of 30% and 60%, respectively (**Fig. 1T**). The speed of protrusion growth versus the speed of retractions was also analyzed using ADAPT software as is explain in the methodology section. It was found that both the control cells and the morphant cells did not show significant differences in the protrusion growth rate. However, the morphant cells exhibited a clear tendency to be slower than the control cells, presenting a rate of 0.89 μm/min ± 0.88 for morphant cells compared to 2.06 μm/min ± 0.51 for control cells (**Fig. 2S**). Moreover, morphant cells exhibited a significant decrease in retraction speed (0.26 μm/min ± 0.13) compared with control cells (2.04 μm/min ± 0.72) (**Fig. 1U**), suggesting that Gαi2 could be influencing the cellular machinery involved in cell body retraction during migration. Therefore, these results collectively, indicate that Gαi2 is required for NC migration *in vivo* and *in vitro*, and its loss of function generates longer cell with bigger protrusions.

**Figure 2.**
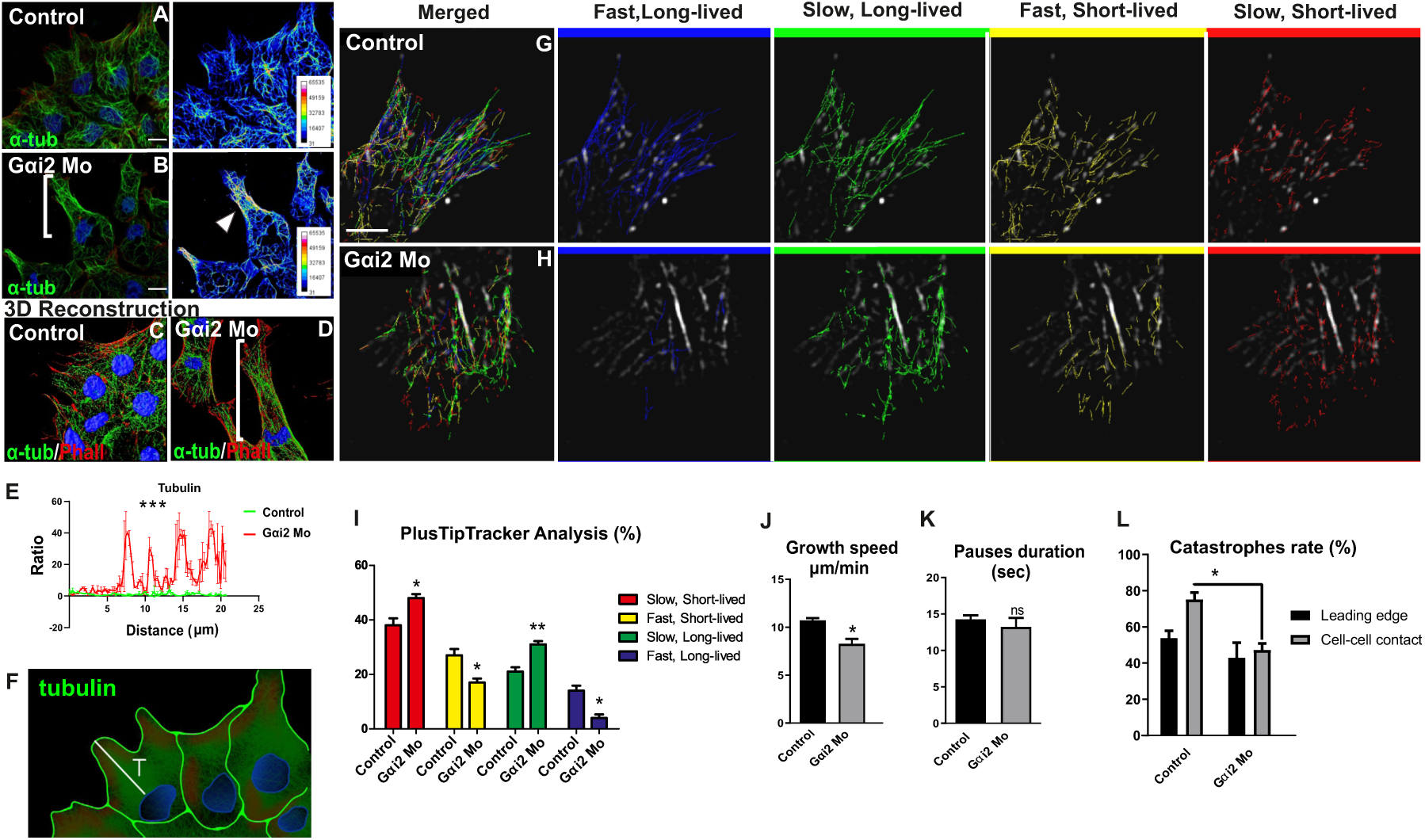
Gαi2 knockdown increases microtubules stability. **A-B.** Immunofluorescence against α-tubulin in *X.t*. Pseudocolor scale shows that Gαi2MO increases microtubules distribution towards the leading edge. Scale bar: 10 μm. Magnification: 63X. **C-D.** Tubulin 3D reconstruction in control and Gαi2 knockdown conditions. **E.** Quantification of tubulin fluorescence intensity. Gαi2 dramatically increased tubulin signal towards the leading edge. Error bars: s.e.m. *** P≤0.001. n= 3 cells were analyzed per condition for each of three independent experiments. **F.** Schematic representation of the tubulin distribution measurements from the edge of the nucleus to the cell cortex. T: tubulin. **G-H.** Images were obtained every 1.5 seconds for a total of 5 minutes using Leica SP8 confocal microscopy. Videos of EB3GFP-labeled cells from *Xenopus laevis* (*X.l*) were cropped to a region of interest, and they were analyzed using plusTipTracker software to detect microtubule growing tips via +TIP tracking proteins (Applegate et al., 2011). Magnification: 63X. **I.** Cells were separated into four groups based on growth rate (slow < 11 µm/min < fast) and duration (short < 23 sec < long). Cells injected with Gαi2MO present a higher percentage of slow microtubules, and a lower percentage of fast microtubules compared to control cells. **J.** The average growth rate of Gαi2 morphant microtubules is 8.52 µm/min ± 0.301 and is significantly lower compared to control cells, whose growth rate is on average 10.53 µm/min ± 0.297. ANOVA statistical analysis for multiple conditions and Student’s T-Test (two-tailed) were performed. N = 3 experiments per condition. **K.** The quantification of the arrests (pauses) of the comets did not show significant differences between control cells and morphant cells for Gαi2. **L.** Using TrackMate plugin of ImageJ, the trajectory of comets was tracked in time, specifically those that enter or leave a pre-established region of interest at the cell-cell contact and at the leading edge. At the end of the 2.5 minutes, a relationship was made between the comets that remain in the area versus the total comets that entered the area. Control cells show a 53.8% of catastrophes at the leading edge and 75.1% at cell-cell contact. On the other hand, Gαi2 morphant cells show 42.9% of catastrophes at the leading edge versus only 47.3% at the cell-cell contact.

### Gαi2 regulates microtubules dynamic during migration

To further explore the potential molecular mechanism by which Gαi2 regulates the morphology and migration of cranial NC cells, we proceeded with the analysis of microtubules cytoskeletal components. With the Gαi/o antecedents about its control on microtubules array during asymmetric cell division (Wang et al., 2005; Kiyomitsu T, 2019; Schaefer et al., 2001; Yu et al., 2000; Villaseca et al., 2022), we first analyzed microtubules cytoskeletal organization through immunofluorescence against α-tubulin. Our findings reveal that microtubules from control cells exhibit a radial arrangement extending towards the leading edge (**Fig. 2A,C).** Conversely, besides an elongated shape, Gαi2 morphant cells exhibit an aberrant cytoskeleton characterized by concentrated distribution of microtubules towards the leading edge **(Fig. 2B,D).** To enhance visualization, we conducted a 3D deconvolution (**Fig, 2C,D, supplementary material movie S2).** This observation is also further supported by the fluorescent intensity quantification graphs, which depict that Gαi2 morphant cells display a clear increase in the tubulin fluorescens signal toward the leading edge (**Fig. 2E,F**).

To further characterize the effect of Gαi2 on microtubules organization we injected mRNA encoding EB3-GFP, a GFP-tagged protein that binds to the plus-end of microtubules. Fluorescent comets representing the growing microtubules were observed using confocal microscope live-cell imaging. We conducted time-lapse tracking of EB3-GFP comets at both 3-second and 1.5-second intervals and quantified various parameters, including growth speed, growth length, comet lifetime, and pauses, using automated analysis with the PlusTipTracker software for Matlab (Applegate et al., 2011; Moore et al., 2013). Comets were classified as either slow or fast and short-lived or long-lived, determined by the mean growth velocity (11 μm/minute) and comet lifetime (23 seconds). Inhibition of Gαi2 expression reduced the proportion of fast, short-lived microtubules in favor of slow microtubules of both short and long-lived (**Fig. 2G,H,I, supplementary material movie S3**). Consequently, microtubules in Gαi2 morphant cells grow more slowly (8.52 μm/min) than those in control cells (10.53 μm/min) (**Fig. 2J**). These results indicate that microtubules originating from Gαi2 morphant cells are less dynamic. Importantly, no significant differences were observed between control cells and Gαi2 morphant cells in the pauses rate experienced by EB3-GFP labeled comets (**Fig. 2K**). On the other hand, the percentage of microtubules catastrophes was calculated using the MTrackJ plugin in ImageJ. For this purpose, we tracked EB3-GFP comets and compared the number of microtubules undergoing catastrophes (disappearing comets) with the total number of microtubules (comets) at the leading edge and cell-cell contact. In control cells, microtubules at the cell-cell contact experienced a higher percentage of catastrophes compared to microtubules at the leading edge (**Fig. 2L**). However, upon Gαi2 knockdown, the percentage of catastrophes, although similar between leading edge and cell-cell contact, was significantly lower at the cell-cell contact compared to control cells (**Fig. 2L**). This suggests that Gαi2 is required to promote microtubules catastrophe during the migration of *Xenopus* cranial NC cells, which correlates with a decrease in microtubules dynamics. Therefore, Gαi2 knockdown leads to increased microtubules stability, resulting in reduced dynamic behavior and a decreased incidence of catastrophes when compared to the control cell population.

### Gαi2 controls Rac1-GTP polarity during cranial neural crest cell migration via microtubule dynamics

Microtubules and actin filaments interact via Rho family of GTPases (Rac1, Cdc42, RhoA), controlling cell polarity, actin polymerization, and contractility (Rodriguez et al., 2003). Microtubules play a crucial role in cell migration, affecting cellular trafficking, morphology, and polarity (Etienne-Manneville, 2013). Altered microtubule polymerization inhibits Rac1 activity, impacting lamellipodium formation (Laan et al., 2008; Waterman-Storer et al., 1999). Several studies have proposed that Gαi is involved in regulating mitotic spindle dynamics, thereby influencing asymmetric cell division and cellular polarity (Kiyomtsu T, 2019; Schaefer et al., 2001; Yu et al., 2000; Villaseca et al., 2022), which is turn is known to be regulated by the Rho family GTPases (Gotta et al., 2001; Cowan and Hyman, 2007). In this study, we observed that Gαi2 morphant cranial NC cells exhibit impaired migration and abnormal cell morphology. These behaviors are closely associated with the organization of the actin and microtubules cytoskeletons, processes finely tuning by the Rho GTPases family proteins. Therefore, to delve deeper into the functional mechanism of Gαi2 in cranial NC cells migration, we aimed to determine whether Gαi2 plays a crucial role in establishing proper cellular polarization during cell migration. To achieve this, we explored whether Gαi2 regulates the subcellular distribution of active Rac1 and RhoA in cranial NC explants under Gαi2 loss-of-function conditions, considering Rho GTPases pivotal roles in cranial NC migration and contact inhibition of locomotion (CIL) (Carmona-Fontaine et al., 2011; Moore et al., 2013; Leal et al., 2018). Hence, we employed mRNA encoding the small GTPase-based probe, enabling specific visualization of the GTP-bound states of these proteins. Cranial NC explants were cultured on fibronectin, and the dynamic localization of the active GTPase was monitored using live-cell confocal microscopy. Consistent with earlier observations by Carmona-Fontaine et al. (2011), in control cranial NC cells, active Rac1 displayed prominent localization at the leading edge of migrating cells, whereas its presence was reduced at cell-cell contacts, coincident with an increase in RhoA-GTP levels (white arrows in **Fig. 3A, supplementary material Figure S3A,C**). On the contrary, in comparison to the control cells, Gαi2 morphants exhibit a pronounced accumulation of active Rac1 protein in the protrusions and at cell-cell contacts, where active RhoA localization is conventionally expected (white arrow in **Fig. 3B, supplementary material Figure S3A,C and movie S4**). In contrast to control cells, a notable shift in the localization of active RhoA protein was observed, with its predominant accumulation now detected at the leading edge of the cell, instead of the typical localization towards the trailing edge or cell-cell contacts (**supplementary material Figure S3B,C).** These findings suggest a dysregulation of contractile forces that align with the observed distribution of active RhoA, cortical actin disruption, and diminished retraction in cell treated with Gαi2MO. As microtubules polymerization locally activates Rac1, resulting in an increase in Rac1 activity towards the protrusions (Waterman-Storer et al., 1999), we aimed to explore whether the microtubules depolymerizing drug nocodazole could reduce the levels of active Rac1 in Gαi2 morphant cells. As demonstrated by the results in (**Fig. 3C-F)**, the administration of nocodazole (65 nM) diminished the levels of active Rac1 in Gαi2 morphant cells. Additionally, immunofluorescence assays showed a change at the localization, accompanied by reduced fluorescence intensity, of both polarity markers, Par3 and ζPKC (**supplementary material Fig. S3D-G)** suggesting a severe impact on cellular polarity under diminished Gαi2 expression.

**Figure 3.**
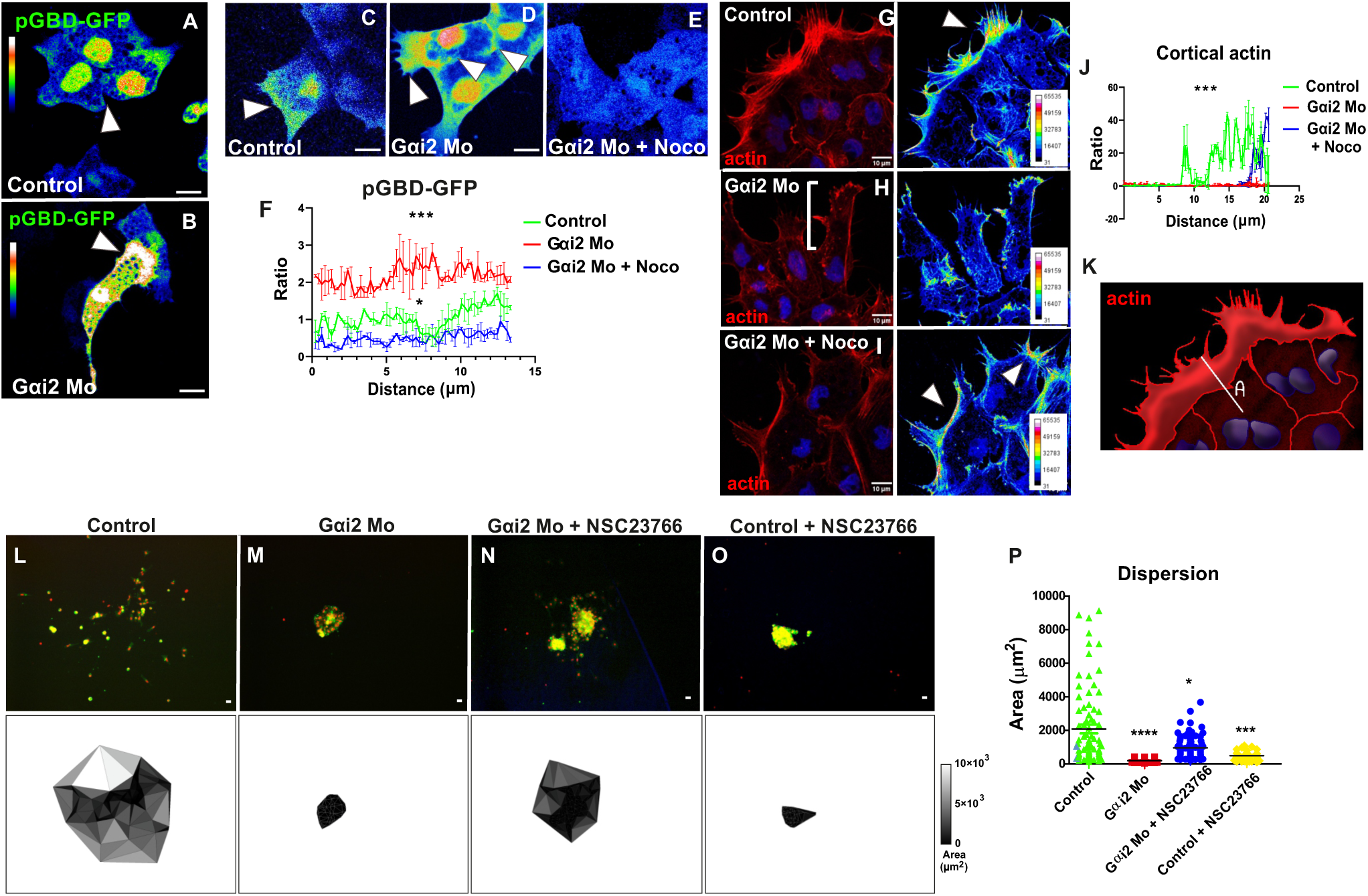
Gαi2 inhibits Rac1 activity in the protrusions and actin turnover via microtubules dynamic. **A-B.** Pseudocolor scale for visualization of changes in the active Rac1 localization via pGBD-GFP probe. This probe contains the Rac1 binding domain of the effector protein PAK1 fused to GFP, which makes it possible to observe the localization of active Rac1 over time. In control cells from *X.t*, active Rac1 is located towards the leading edge and its presence is reduced at the cell-cell contact (white arrow). In Gαi2 morphant cells from *X.t*, active Rac1 is localized throughout the cell, and highly concentrated in the cell protrusion at the cell-cell contact (white arrow). Scale bar: 10 μm. **C-E.** Gαi2 morphant cells from *X.t* treated with nocodazole show a decrease in the levels of active Rac1. Scale bar: 10 μm. **F.** Quantification of active Rac1 fluorescence intensity in time. The significance was evaluated with a Mann-Whitney test for non-parametric data (***, p<0.001; *, p<0.05). Error bars: s.e.m. Magnification: 40X. (experimental N = 3). **G-I.** Actin was stained with phalloidin to visualize cortical actin in Control, Gαi2 morphant condition and Gαi2 knockdown condition under 65 nM nocodazole treatment. Scale bar: 10 μm. **J.** Quantification of actin fluorescence intensity showing that Gαi2 MO severely affects cortical actin distribution. Error Bars: s.e.m. *** P≤0.001, * P≤0.05. n= 3 cells were analyzed per condition for 3 each independent experiments. **K.** Schematic representation of the measurements for actin distribution specifically from the edge of the nucleus to the cell cortex. A: actin. **L-O.** Cranial NC explants were cultured into fibronectin and cell dispersion was evaluated during 4 hours through timelapse. All embryos were co-injected with H2B-Cherry and membrane GFP to monitor individual cells and quantify cell dispersion. Four conditions were evaluated: Control, Gαi2MO, Gαi2MO + 20 nM NSC23766 and Control + 20 nM NSC23766. Delaunay triangulation of a representative explant from *X.t* of each condition at the final time (4 hours) is show below every explant. **P.** The radius between the average areas corresponding to the final time were plotted at 4 hours to evaluate cell dispersion. Significance was assessed with a Mann-Whitney test for non-parametric data (****, P<0.0001). Error bars: s.e.m. 10X magnification. Control n = 15 explants; Gαi2MO n = 18 explants; Gαi2 Mo + 20 nM NSC23766 = 14 explants; Control + 20 nM NSC23766 = 12 explants; 3 trials.

The extension of the plasma membrane to form lamellipodia is primarily driven by actin polymerization, a process mainly regulated by Rac1 (Ridley A., 2015). The concurrent inhibition of active Rac1 at the leading edge significantly impacts cellular polarity and migration (Moore et al., 2013), making any changes in the levels of these GTPases critically influential. To further comprehend the impact of Gαi2 knockdown on Rac1 activity as we found in this work, we proceeded to investigate the potential rescue of cell migration behavior by employing a pharmacological inhibitor (NSC23766) to modulate Rac1 activity. It is worth noting that we conducted Rac inhibitor NSC23766 trials at concentrations ranging from 20 nM to 50 nM for *X. laevis* and between 10 nM to 30 nM for *X. tropicalis*. In both cases, higher concentrations of the Rac inhibitor proved to be lethal (data not shown), underscoring the essential role of Rac1 in both cell migration and cell survival. Remarkably, our results show partial restoration in cranial NC cells dispersion following a 5-minute treatment with a low concentration of the Rac1 inhibitor (20 nM of NSC23766 *X. laevis* and 10 nM for *X. tropicalis*) (**Fig. 3L-P, supplementary material movie S5)**. This suggests that these concentrations are sufficient to demonstrate that the increase in Rac1-GTP resulting from Gαi2 morpholino knockdown impairs cell migration. The Rac1 inhibitor was administered by adding the molecule to the solution, leading to its inhibitory effect being observed both at the leading edge and the cell-cell contacts, significantly disrupting cell migration. This effect was evident in the control cells treated with the Rac1 inhibitor, as they exhibited a complete loss of migratory capacity due to reduced cell dispersion (**Fig. 3O,P, supplementary material movie S5**). This finding suggests that the observed partial rescue of migration may be attributed to the global inhibition of Rac1 throughout the cell, resulting from the method of treatment. Therefore, we believe that the lower concentration of NSC23766 used in our assay was adequate to reduce the abnormal Rac1-GTP activity in the morphant NC cells. However, it is important to note that for normal NC cells, this level of reduction in Rac1-GTP activity is critical and sufficient to impair normal migration. All these findings collectively indicate that Gαi2 plays a pivotal role in the regulation of cell polarization during cranial NC cell migration, possible controlling Rac1 activity via microtubules dynamics. A decrease in Gαi2 levels leads to distinct changes in the subcellular localization of all the analyzed polarity markers. Therefore, these alterations could be potentially be associated to disruptions in the microtubule network which, in turn, may impact the proper transportation of cell polarity proteins.

As we mention above, Rac1 activity controls protrusion formation, consequently we next analyzed actin cytoskeleton organization by immunofluorescence against F-actin. We found that in control cells cortical actin is typically localized at the leading edge establishing normal protrusions (**Fig. 3G**). On the contrary, Gαi2 morphant cells display not only enlarged protrusions, as demonstrated above, but also an aberrant cytoskeleton marked by disorganized cortical actin cables, nearly disappearing at the leading edge (**Fig. 3H**). Therefore, considering that Gαi2 knockdown conditions showed microtubules highly concentrated towards the protrusions at the leading edge (**Fig. 2B**), we proceeded to evaluate whether cortical actin organization can also be rescued with nocodazole. Consistently, the treatment of Gαi2 morphant cells with 10 μM of nocodazole resulted in a partial restoration of the normal organization of cortical actin (**Fig. 3I**), supported by the fluorescence intensity quantification graph (**Fig. 3J,K**). All together these findings suggest that during cranial NC cells migration, Gαi2 inhibits Rac1 activity in the protrusions and probably actin turnover, via microtubules dynamic.

### Gαi2 regulates focal adhesion dynamics by controlling their disassembly during cell migration via microtubules dynamic

Our investigation reveals that Gαi2 morphant cells exhibit noticeable migration defects and substantial alterations in cellular morphology, particularly manifesting a significant increase in protrusion area. Moreover, these cells display distinct changes in the organization of the actin and microtubules cytoskeletons, characterized by a concentrated distribution of microtubules towards the leading edge. Considering the well-established role of microtubules in facilitating the transport of proteins involved in diverse dynamic processes, including focal adhesion dynamics, we are prompted to explore the potential association of these abnormal morphological and cytoskeletal phenotypes with defects in focal adhesion assembly or disassembly. To assess the contribution of Gαi2 in the focal adhesion morphology and dynamic, an immunohistochemical analysis was conducted against β-integrin and phospho-paxillin (p-Pax), both key proteins involved in focal adhesion assembly. It has been reported that integrins, once in their active conformation, recruit several adapter proteins with kinase activity, including FAK, Src, and Paxillin (Parsons et al., 2012). Cells with Gαi2 loss of function exhibited an increase in the area of focal adhesions compared to control cells (**Fig. 4A-D**). Additionally, the dynamics of focal adhesions was evaluated by conducting time-lapse imaging of a GFP-tagged form of FAK expressed in explanted NC cells. Our results revealed a significantly higher stability of focal adhesion in Gαi2 morphant cells compared to control cells (**Fig. 4E-H**, **supplementary material movie S6**). These findings suggest that integrins and the adapter protein FAK comprising the focal adhesions remain localized at the cell edge for an extended duration. These results suggest that Gαi2 morphant cells exhibit a slower dynamic of focal adhesions disassembly, in contrast to control cells, which may be related to the regulatory role of Gαi2 on microtubules organization.

**Figure 4.**
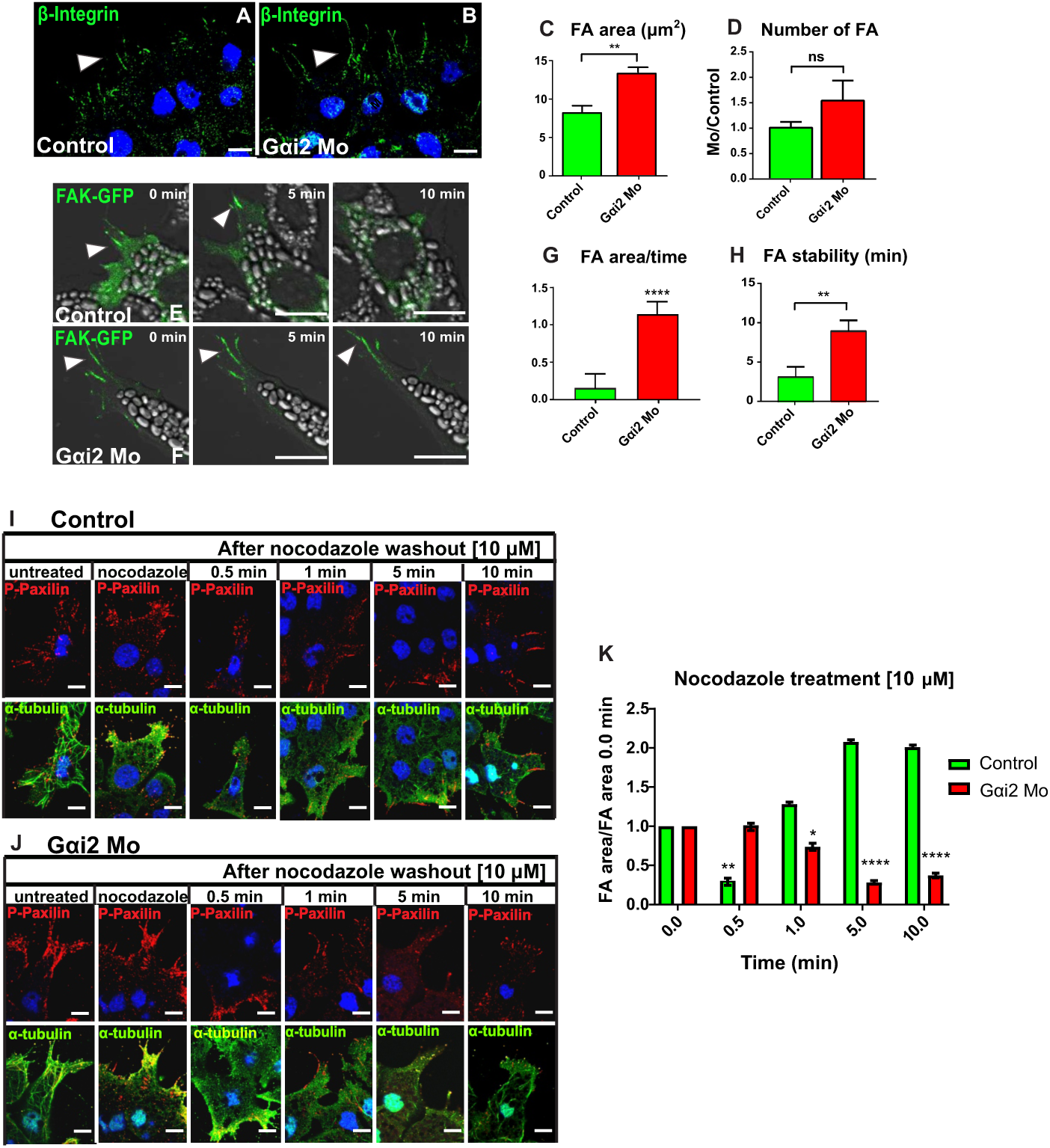
Gαi2 morpholino inhibits focal adhesion turnover via microtubules stability. **A-B.** Immunostaining against β1-integrin in Control and Gαi2 morphant conditions to detect focal adhesions. In *X.t.* Scale bar: 10 μm. **C-D.** Quantification of the area and number of focal adhesions. Statistical analysis using t-student (**, p<0.01; ns: non-significative; n explants = 43). **E-F.** Images obtained by SP8 confocal microscopy timelapse following FAK-GFP localization in control and Gαi2 morphant conditions from *X.t* explants. Scale bar: 10 μm. Magnification: 63X. **G-H.** Quantifications of changes in the focal adhesion area during time and focal adhesion stability. Focal adhesion from Gαi2 morphant cells are more stable over time, compared to control cells. Statistical analysis using t-student (**, P<0.01; ****, P<0.0001; n explants = 11). **I-J.** Immunofluorescence performed to detect endogenous p-Pax (red) and endogenous α-tubulin (green) in control and Gαi2 knockdown conditions from *X.t* explants, under nocodazole treatment (10 µM). Scale bar: 10 μm. **K.** Normalized graph from Gαi2 knockdown condition shows an increase in the focal adhesion area, decreasing the disassembly dynamic of focal adhesion after nocodazole treatment. We can observe that control focal adhesions disassembly starts at 0.5 minutes, however Gαi2 morphant explants show a delay in focal adhesion disassembly, which start 5 minutes after nocodazole wash out. Error bars: s.e.m., * P < 0.05; ** P> 0.01; **** P < 0.0001. n explants = 27, 3 experimental replicates. Magnification: 63X.

Therefore, to evaluate the dynamics of microtubules-dependent focal adhesion assembly and disassembly, both control and Gαi2 morphant conditions, were synchronized using 65 nM of nocodazole to deplete the microtubules network, thereby immobilizing the adhesions on the cell membrane. Subsequently, the cells were washed to restore microtubules polymerization, and facilitate cargo transport, and fixed at various time points to establish a temporal profile of focal adhesions assembly and disassembly dynamics. Immunofluorescence staining was performed against p-Pax and α-tubulin, and the area of focal adhesions was analyzed over time. In control cells, a rapid decrease in focal adhesion area is observed within 30 seconds after the nocodazole washes (**Fig. 4I,K**). Furthermore, a drastic increase in focal adhesion area occurs at one minute (**Fig. 4K**, green bars), indicating a 30-second period for both assembly and disassembly. In contrast, Gαi2 morphant cells exhibit a reduction in focal adhesion area only after 5 minutes, which persists until 10 minutes (**Fig. 4J,K**, red bars), revealing a time frame of approximately 5 minutes for the disassembly process, significantly different from the control cells, which exhibited a brief 30-second between each process. These findings demonstrate that the focal adhesion disassembly process is significantly slower in Gαi2 morphant cells compared to control cells (**Fig. 4K**). Taken together, these data strongly support the conclusion that Gαi2 exerts a regulatory role in the disassembly dynamics of focal adhesions, by controlling microtubules dynamic, resulting in a significant impact on *in vitro* migration.

### Gαi2 interacts with plus end proteins EB to regulate microtubules dynamics

To identify the mechanism by which Gαi2 regulates cranial NC cells migration through the control of microtubules distribution and dynamics, an analysis was conducted to determine which population of microtubules is concentrated in the cell. Our results revealed a significant increase in acetylated tubulin levels, indicative of more stable microtubules in Gαi2 morphant cells, whereas control cells exhibited reduced acetylated tubulin, as evidenced by western blot analysis, which quantification graph display a gray value of approximately 1.5, in morphant cells compare with 0.5 in control cells (**Fig. 5A,B**). This observation was further validated by immunofluorescence, revealing an increase in acetylated tubulin levels within morphant cells, particularly towards the protrusions at the leading edge (**supplementary material Fig. S4C-I**). In addition, the subcellular localization of Gαi2 in relation to α-tubulin was subjected to analysis, demonstrating colocalization of both proteins within the same subcellular compartment (**Fig. 5C-E**). The quantification, using the Pearson coefficient, demonstrated a strong correlation with a value of 0.743, while the overlap coefficient was 0.779 (**Fig. 5F**). These results are supported through co-immunoprecipitation and proximity ligation assays (PLA). The co-immunoprecipitations show that Gαi2 and tubulin are part of a common interaction complex (**Fig. 5G, supplementary material Fig. S4B**) along with microtubules plus-end proteins, such as EB3 **(Fig. 5H, supplementary material Fig. S4B**) and EB1 (**Fig. 5I**), which are associated with tubulin to regulate microtubule dynamics (Perez et al., 1999; Bieling et al., 2007). A V5 antibody alone was utilized as a control (**supplementary material Fig. S4A).** Notably, supporting these results, PLA demonstrates a direct interaction of Gαi2 with EB1 but not directly with tubulin *ex vivo* and *in situ*, as indicated by comet-dots type imaging directed towards the leading edge of NC cells (**Fig 5J-M**). PLA utilizing specific antibodies against EB1 and α-tubulin, as a control, exhibited robust and highly specific signals in the form of multiple cytoplasmic dots towards the leading edge (**Fig. 5J,N**). In contrast, Gαi2 and α-tubulin did not show interaction by PLA, potentially due to both proteins being components of the same complex, as demonstrated in the Co-IPP (**Fig. 5G**), although with a higher distance of 40 nm between them (**Fig. 5K**). However, Gαi2 and EB1 exhibited a similar pattern to EB1 and α-tubulin, with specific signals observed as multiple cytoplasmic dots towards the leading edge, albeit with a decreased number of dots (**Fig. 5L,N**), possibly indicating a decreased number of interactions. Interestingly, the number of dots in both cases, Gαi2-EB1 interaction, and EB1-α-tubulin interaction, was higher in the outer cells of the explants compared to the inner cells (**Fig. 5O**), suggesting a greater degree cytoskeletal microtubule organization towards the direction of migration. Therefore, these findings suggest a functional and physical relationship between Gαi2 and plus-end binding proteins, such as EB1, and through EB1 with tubulin, thus regulating microtubule dynamics. The significant alterations in tubulin organization observed under Gαi2 morphant conditions and the increased presence of stable tubulin at the leading edge further support this notion.

**Figure 5.**
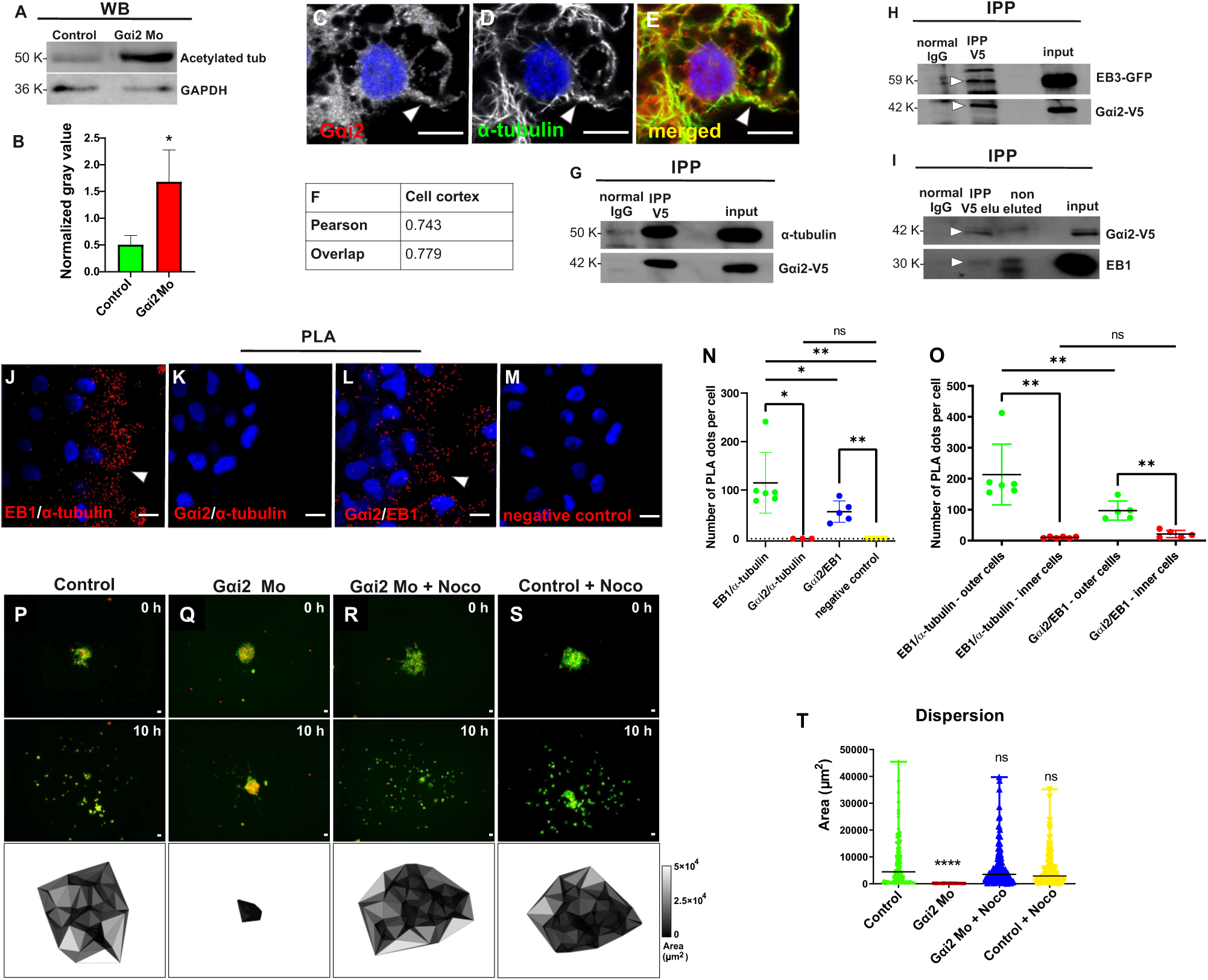
Gαi2 interacts with microtubules through the plus tip end protein EB1, controlling microtubules dynamics and therefore cranial NC cells migration. **A-B.** Western blot performed to measure the quantity of acetylated tubulin under control and Gαi2 knockdown conditions. Intensity was normalized to GAPDH to quantifying the increase in the acetylated tubulin amount in morphant conditions. **C-E.** Immunofluorescence against endogenous Gαi2 and α-tubulin proteins to study subcellular localization. Scale bar: 10 μm. **F.** Protein colocalization was analyzed using Image’s JACoP software whose Pearson coefficient was 0.743, while the overlap coefficient was 0.779. **G.** Coimmunoprecipitation assays show that Gαi2 is interacting with α-tubulin. normal IgG: non-related IgG; IPP V5: immunoprecipitation conjugated to V5 antibody; input: total lysate. n total = 3 lysates per condition. **H-I.** Coimmunoprecipitation assays show that Gαi2 interacts with EB1 and EB3, plusTip proteins that regulate microtubules dynamic. W/A: Without antibody; normal IgG: non-related IgG; IPP V5: immunoprecipitation conjugated to V5 antibody; input: total lysate; elu: eluted fraction. Non-eluted: non-eluted fraction. n total = 3 lysates per condition. White arrows highlight the correct protein bands. **J-M.** Proximity Ligation Assay (PLA) performed for *in situ* detection of Gαi2 interaction with plus end proteins, unveiling that Gαi2 interacts with tubulin through EB1. White arrows indicate the interaction detected at the leading edge. EB1/α-tubulin: positive control, negative control: without primary antibody. Scale bar: 10 μm. **N-O.** Quantification of PLA Dots per Cell in Outer and Inner Cells. The graphs illustrate the mean number of PLA dots per cell for the entire cell explant population **(N)**, as well as separately for outer and inner cells **(O)**. The graphs depict the mean number of PLA dots per cell, with statistical analysis conducted using the Mann-Whitney test for non-parametric data (**P < 0.05, ns: non-significative;). Error bars: s.e.m. **P-S.** Cranial NC explants were extracted from stage 17 *X.l.* embryos, previously injected with Gαi2MO. The cell dispersion in each condition was evaluated during 10 hours through timelapse. All embryos were co-injected with H2B-Cherry and membrane GFP for individual cell monitoring and quantification of migration. Four conditions were evaluated: Control, Gαi2MO, Gαi2MO + 65 nM Nocodazole and Control + 65 nM Nocodazole. Under each representative image, Delaunay triangulation is shown in each condition at the final time (10 hours). **T.** The rate between the average areas corresponding to the final time were plotted at 10 hours. Nocodazole can rescue Gαi2 morphant phenotype and does not affect control migration. Significance was assessed with a Mann-Whitney test for non-parametric data (****, P<0.0001). Error bars: s.e.m. 10X magnification. Control n = 21 explants; Gαi2MO n = 24 explants; Gαi2MO + 65 nM Nocodazole n = 36 explants; Control + 65 nM Nocodazole n = 30 explants; 3 experiments.

For a more functional insight into the role of Gαi2 in regulating cranial NC cells migration through the control of microtubules dynamics, we conducted a rescue experiment using a low concentration of nocodazole (65 nM) and cell dispersion *in vitro* was analyzed by Delaunay triangulation. Our results indicated that the treatment with this concentration of nocodazole had no significant impact on the dispersion of control cells (**Fig. 5P,S,T, supplementary material movie S7**). However, in the context of Gαi2 morphant cells, where we observed that Gαi2MO blocked cranial NC cells dispersion (**Fig. 5Q,T**), treatment with 65 nM of nocodazole efficiently rescued cell dispersion (**Fig. 5R,T, supplementary material movie S7**) and additionally cell morphology (**supplementary material movie S8)**, suggesting that the loss of function of Gαi2 increases microtubules stability, phenotype that can be rescued by microtubule depolymerization using nocodazole.

All together, these findings further support the notion that Gαi2 control cranial NC cell morphology and migration regulating microtubules dynamics, by interacting with microtubules plusTip proteins such as EB1 and EB3.

Collectively, our results provide compelling evidence that Gαi2 controls microtubule dynamics by interacting with accessory proteins such as EB1 and EB3, therefore regulating cell morphology and cell-matrix adhesion through the involvement of the small GTPase, Rac1. This regulatory mechanism establishes a novel crucial coordination and crosstalk between the actin and tubulin cytoskeletons and Gαi2 subunit from heterotrimeric G protein, essential for the orchestrated collective migration of cranial neural crest cells.

## Discussion

For effective migration, dynamic microtubules are crucial. They act as transport routes for proteins, facilitating the delivery and recycle of membrane components and signaling molecules to the leading edge of migrating cells (Bershadsky et al., 1996; Bouchet and Akhmanova, 2017; Etienne-Manneville, 2013; Garcin and Straube, 2019). Gα subunits of heterotrimeric G proteins are key regulators of migration in various cell types (Cotton and Claing, 2009), with Gαi subunits playing pivotal roles in immune cell migration (Hwang et al., 2007; Wiege et al., 2012; Zhong et al., 2012; Hwang et al., 2015). In this study, we demonstrate that the functional deficiency of Gαi2, a protein we previously identified as having significant expression in *Xenopus* cranial NC cells and cranial NC-derived cells (Fuentealba et al., 2016), does not impair NC induction and the transition from E-cadherin to N-cadherin expression. As a result, the primary role of Gαi2 become evident during the dynamic process of active NC migration. We made a significant finding that the Gαi2 subunit of heterotrimeric G proteins plays a pivotal role in regulating microtubule dynamics by interacting with their regulatory proteins, such as EB1 and EB3, during microtubules assembly and disassembly dynamics. This regulation, in turn, influences cellular polarity components, including Rac1 and RhoA GTPases, and has a direct impact on actin polymerization and, consequently, the dynamics of focal adhesions (**Fig. 6**). These cellular processes are crucial in governing cell migration.

**Figure 6.**
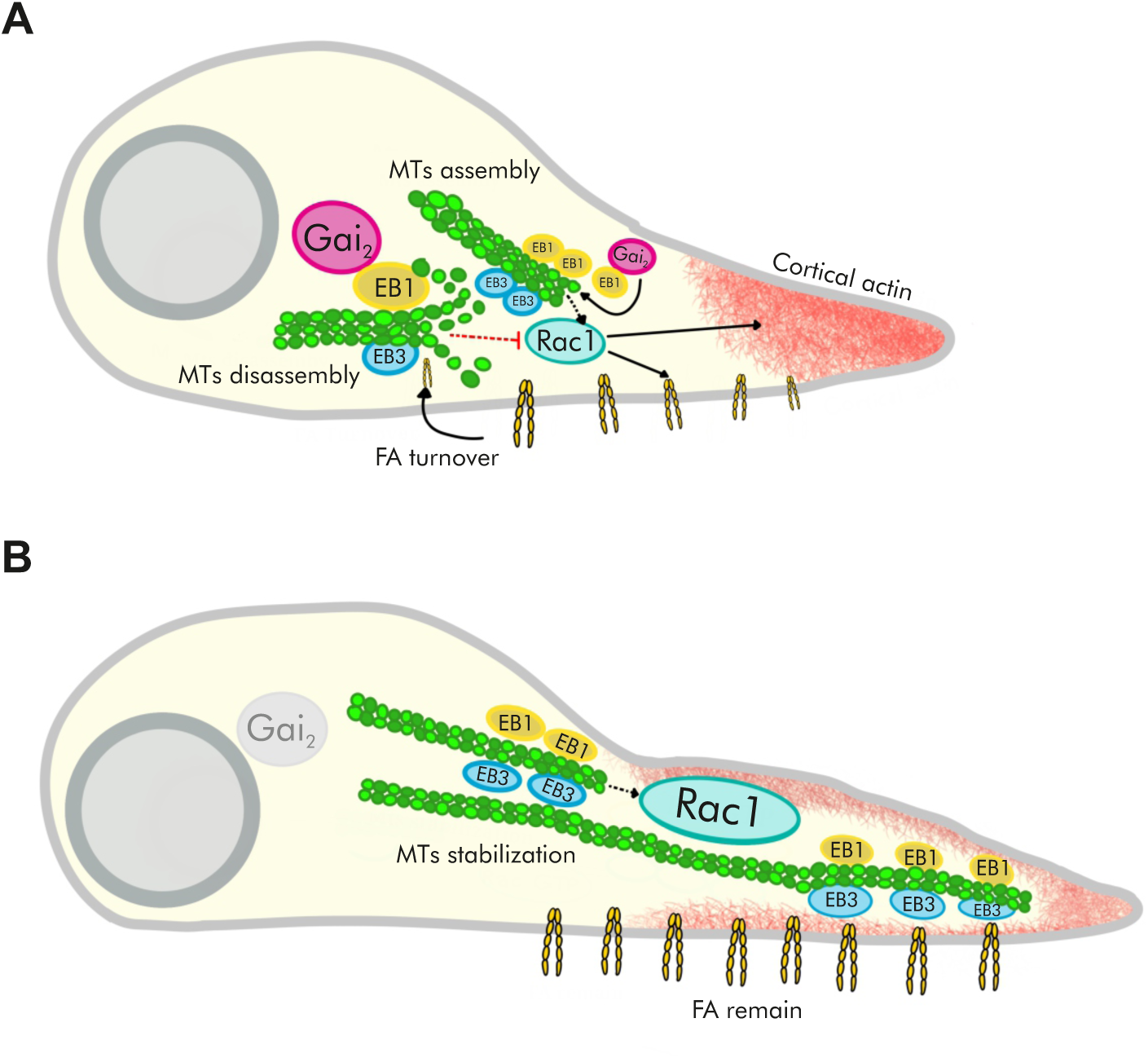
Gαi2 controls cranial NC migration by interacting with EB1 regulating microtubules dynamics. **A.** In the control condition, Gαi2-expressing cells maintain dynamic microtubules by the Gαi2 binding EB1 to the plus end of microtubules. These dynamic microtubule networks control Rac1 activity at the leading edge, possible by intermediary proteins, such as GDIs, GEFs and GAPs, promoting cortical actin formation in the leading cells, facilitating rapid focal adhesion disassembly and enabling a change in the direction of migration. **B.** Conversely, in Gαi2 knockdown conditions, cells have reduced microtubules dynamics, a decreased catastrophe rate, stable and larger focal adhesions, and slower and more stable microtubules wherein EB3 appears to remain persistently associated with them and increasing tubulin acetylation. Stable microtubules lead to enhanced Rac1 activation at the leading edge, interfering with cortical actin formation and FA disassembly, thus inhibiting collective migration.

Our initial findings reveal that the functional loss of Gαi2 significantly inhibits both *in vivo* and *in vitro* migration. Interestingly, during our *in vitro* dispersion assays on fibronectin, control cells displayed radial dispersion, facilitated by the initial contact inhibition force, which allowed them to separate from the group and migrate. In contrast, Gαi2 morphant cells exhibited reduced dispersion efficiency, showing a tendency to remain within the cellular cluster. This cellular behavior in the knockdown condition of Gαi2 is consistent with the observations made by Wiege et al. in 2012, where they reported that Gαi2-/- mouse-derived macrophages lost their migratory capacity towards inflamed regions in alveolitis and peritonitis cases, indicating a compromised cellular motility. Therefore, in order to gain deeper insights into the mechanism underlying Gαi2-mediated regulation of cellular migration, we conducted an analysis of cellular morphology, with a specific focus on actin dynamics, in both the control and Gαi2 morphant conditions. As observed in our results, the Gαi2 morphant condition exhibited a statistically significant increase in protrusion area compared to control cells, whereas the retraction area was considerably reduced in the Gαi2 morphant condition. Additionally, our assessment of retraction velocity demonstrated a significant decrease in Gαi2 morphant cells, suggesting a prolonged stability of protrusions over time compared to control cells, which exhibited minimal retraction and tended to display larger protrusion areas. These findings implicate Gαi2 in potential signaling pathways involved in the regulation of protrusion formation thereby impacting the dynamic of cell migration.

This notion is consistent with our examination of tubulin distribution that unveiled a distinctive pattern in Gαi2 morphant cells, with microtubules concentrated towards the leading edge in parallel bundles, in contrast to control cells exhibiting radial tubulin distribution. This suggests that Gαi2 knockdown provoke changes in the microtubule dynamics towards the leading edge, potentially interfering with cortical actin formation in this region. Gαi proteins regulate astral microtubule organization during asymmetric cell division (Yu et al., 2000; Parmentier et al., 2000; Gotta and Ahringer, 2001; Schaefer et al., 2001; Villaseca et al., 2022) probably by modulating microtubule dynamics. In their absence, microtubule catastrophes decrease *in vitro*, as Gαi inhibits assembly and activates tubulin intrinsic GTPase, enhancing microtubule dynamics by removing the GTP cap by interacting with intermediary proteins, such as GDI (Roychowdhury et al., 1999; Roychowdhury and Rasenick, 2008). Nevertheless, in this study, we present, for the first time, a novel role for the Gαi2 subunit in controlling microtubule dynamics by intercating with EB1 during cell migration events. This occurs at a point when cell division has ceased, unveiling a new non-canonical role for Gαi2. Our specific analysis of microtubule dynamics in Gαi2 morphant cells, using the GFP-tagged microtubule plus-end tracking protein EB3, revealed a remarkable increase in microtubule stability over time. The growth rate of microtubule in these cells was slower compared to control cells, and the frequency of microtubule catastrophes was reduced, showing clear cell migration defects. Interestingly, in addition to microtubules prominent concentration at the leading edge, they exhibit a compelling phenotype wherein EB3 appears to remain persistently associated with them, resulting in a distinctive long-bundle morphology as opposed to the characteristic comet-like appearance in control cells. Furthermore, our investigation revealed that Gαi2 forms an interaction complex with α-tubulin by directly interacting with the plus-end-binding protein of microtubules, such as EB1, known to regulate microtubule dynamics (Mimori-Kiyosue et al., 2000; Akhmanova and Steinmetz, 2015; Serre et al., 2019). Interestingly, PLA dots, evidencing interaction, are localized towards the leading edge and displaying a higher abundance in the outer cells compared with the inner cells, suggesting an increased cytoskeletal microtubule organization towards the direction of migration. Consistent with these results, when examining the population of microtubules primarily concentrated at the leading edge, we observed a higher percentage of acetylated microtubules in Gαi2 morphant cells compared to control cells. Acetylated microtubules are known to exhibit greater stability (Eshun-Wilson et al., 2019). Earlier investigations have demonstrated that cranial NC cells can be prompted to migrate by reducing microtubules acetylation, even in soft matrices (Marchant et al., 2022). Additionally, research has indicated that modulations in microtubule stability are essential for establishing a polarized microtubule cytoskeleton in motile cells (Garcin and Straube, 2019). Supporting these, a recent study has proposed that the acetyltransferase MEC-17 directly promotes α-tubulin acetylation, leading to the inhibition of cell motility by increasing cell spreading area (Lee et al., 2018). Our findings align with this notion, as we observed that Gαi2 morphant cells exhibit a migration-inhibited phenotype displaying and elongated protrusion area in the presence of highly acetylated and concentrated microtubules at the leading edge. These findings strongly support that Gαi2 plays a role in modulating microtubule dynamics as part of its interaction with regulatory proteins, such as EB1 and EB3.

During cell migration, microtubules extend to the periphery, and their dynamics are influenced by local factors, thereby impacting actin polymerization pathways (Akhshi et al., 2013). Microtubules and actin microfilaments work together to meet cellular demands (Etienne-Manneville et al., 2013; Dogterom et al., 2019; Garcin et al., 2019; Seetharaman et al., 2020). Microtubules participate in protein trafficking that regulates actin polymerization, including EB1 and mDia1-2, which control actin filament assembly from microtubules plus-ends (Henty-Ridilla et al., 2016). Some studies show that as microtubules grow toward the migratory edge, plus-end-binding proteins like CLIP170 enhance their stabilization (Galjart et al., 2010) and closely associate with mDia1, promoting actin polymerization from microtubules plus-ends, contributing to lamellipodia and filopodia formation (Henty-Ridilla et al., 2016). Our findings support this idea, as we observed a complete restoration of the migration *in vitro* and partial restoration of cortical actin and cell morphology in Gαi2 morphants using the microtubule-depolymerizing drug nocodazole. Disassembling the concentrated microtubules at the leading edge indicated that Gαi2 knockdown induces significant changes in microtubule dynamics, leading to their occupation into the cortical actin domains. This, in turn, disrupts actin polymerization, which can be partially rescued with nocodazole.

GTPase Rac1 plays a vital role in actin cytoskeleton remodeling, particularly at the leading edge during cell migration (Nobes et al., 1995; Ridley et al., 2015). Therefore, Gαi2 knockdown is expected to increase active Rac1 at the leading edge, a behavior confirmed in this study and consistent with our findings of increased protrusion areas upon Gαi2 knockdown condition. Previous studies, had linked Gαi2 to CXCR4 and Elmo1/Dock180 complex suggesting a potential molecular interplay with Rac1, promoting its activation (Li et al., 2013). Additionally, the known association of Gαi signaling with Src kinase in ovarian cancer cells activating Rac1 (Ward et al., 2015), provides insights into observed cellular morphology changes in our study. Interestingly, our analysis of the cell cortex revealed that Gαi2 knockdown results in a paradoxical phenomenon: cortical actin is diminished, despite the presence of more stable protrusions. This contradicts the expected increase in active Rac1 and consequent actin polymerization. Other studies have reported that microtubule assembly promotes Rac1 signaling at the leading edge, while microtubule depolymerization stimulates RhoA signaling through guanine nucleotide exchange factors (GEF) associated with microtubule-binding proteins controlling cell contractility, via Rho-ROCK and focal adhesion formation (Krendel et al., 2002; Ren et al., 1999; Best et al., 1996; Garcin and Straube, 2019; Waterman-Storer et al., 1999; Bershadsky et al., 1996; Moore et al., 2013). This mechanism would contribute to establishing the antero-posterior polarity of cells, crucial for maintaining migration directionality, underscoring the significance of regulating microtubule dynamics in directed cell migration. These findings closely align with the results obtained in this investigation, demonstrating that Gαi2 knockdown reduces microtubule catastrophes and promotes tubulin stabilization, resulting in increased localization of active Rac1 at the leading edge and cell-cell contacts, while decreasing active RhoA at the cell-cell contact but increasing it at the leading edge. This possibly reinforces focal adhesion, which is consistent with the presence of large and highly stable focal adhesions under Gαi2 knockdown conditions. This finding also suggests a dysregulation of contractile forces in comparison to control cells, a result that aligns with the observed distribution of active RhoA, cortical actin distribution and diminished retraction in cells treated with Gαi2MO. This strikingly contrasts with the normal cranial NC migration phenotype, where Rac1 is suppressed at cell-cell contacts during CIL, leading to a shift in polarity towards the cell-free edge to sustain directed migration (Theveneau et al., 2010; Shoval and Kalcheim, 2012; Leal et al., 2018). In addition, our findings here, where we observed a partial rescue at the Gαi2 morphant migratory behavior using a low concentration of Rac1 inhibitor (NSC23766) strongly support this possible mechanism. Some studies in epithelial cells have associated Par3 and ζPKC with cell-cell adhesion complexes, crucial for apical-basal cell polarity, and like Gαi in asymmetric cell division (Hao et al., 2010). Par3 is also implicated in regulating microtubule dynamics and Rac1 activation through interaction with Rac-GEF Tiam1 (Chen et al., 2005). Additionally, Par3 has been found to regulate CIL in *Xenopus* cranial NC cells, promoting microtubule catastrophe and inhibiting Rac1/Trio signaling during cell collision (Moore et al., 2013). These results support our findings as Gαi2 morphant cells showed altered subcellular localization of Par3 and ζPKC, possible due to impaired microtubule dynamics, leading to a delayed delivery of these proteins to their required sites. Collectively, all these findings suggest that Gαi2 inhibit Rac1 activity in the protrusions and probably actin turnover, via microtubules dynamic.

Cell migration relies on the detection of various signaling cues, including chemical gradients (chemotaxis), adhesion gradients (haptotaxis), and extracellular matrix (ECM) stiffness gradients (durotaxis) (Piacentino et al., 2020). Microtubules stabilize ECM receptors, called integrins, which form focal adhesions for cell to anchor to the substrate and generate traction forces (Shacks et al., 2019; Rottner et al., 2019; Dogterom et al., 2019; Kaverina et al., 1999). Microtubules also facilitate trafficking of integrins, chemokine receptors, and essential adaptor proteins to regulate focal adhesion dynamics, including Calpain-2 (Etienne-Manneville et al., 2013; Garcin and Straube, 2019; Dogterom et al., 2019; Seetharaman et al., 2020). It has been reported that plus-end microtubule-binding proteins, such as EB1, interact with focal adhesion adaptor proteins, including talin, focal adhesion kinase (FAK), Paxillin, and Src (Bouchet et al., 2016; Dubois et al., 2017). Interestingly, our findings demonstrate a direct interaction between Gαi2 and EB1 revealing that the Gαi2 subunit positively regulates microtubule dynamics, a regulation likely contributing to the maintenance of focal adhesion dynamics during cell migration. FAK, activated by Src-induced phosphorylation, regulates focal adhesion size by recruiting growth factor receptor-binding protein 2 (Grb2) and dynamin for integrin internalization (Mitra et al., 2005; Ezratty et al., 2005). Interestingly, Gαi may impact this process, as it is known to interact with PI3K and Grb2 in response to growth factors like EGF (Cao et al., 2009; Zhang et al., 2015; Li et al., 2016; Liu et al., 2018; Nusková et al., 2021; Villaseca et al., 2022). This suggests that in the Gαi knockdown condition, Grb2 might be unable to recruit dynamin, potentially hindering integrin internalization and reducing focal adhesion size. Therefore, investigating dynamin localization and integrin trafficking in Gαi2 knockdown conditions could provide valuable insights. In our investigation, Gαi2 knockdown cells exhibited significantly reduced focal adhesion disassembly dynamics a process that dependent of microtubules dynamics. This is consistent with previous findings in human fibroblasts and endothelial cells treated with pertussis toxin (PTX), an uncoupler of Gαi subunit from its receptor (Orr et al., 2002; Orr et al., 2003). Additionally, treatment with membrane-permeable peptides blocking Gαi2 subunit function, effectively hindered thrombospondin (TSP)-mediated focal adhesion disassembly. TSP, a matricellular protein that activates PI3K to promote the disassembly of stress fibers and focal adhesions (Orr et al., 2002), has also been studied in chicken embryo NC cells, where it was found to enhance migration speed (Tucker et al., 1999).

It is well established that the activity from the Rho family of small GTPases is controlling cytoskeletal organization during migration (Ridley AJ., 2015). Contrariwise, it has been described in many cell types, that microtubules dynamic polymerization plays a crucial role in establishing the structural foundation for cell polarization, consequently influencing the direction of cell motility (Watanabe et al., 2005). Our results appear to align with this latter view. While it is reasonable to postulate the possibility that Gαi2 regulates Rac1 activity, subsequently influencing actin and microtubule dynamics, our findings in the context of cranial NC cells, lend support to an alternative sequence of events. Initially, Gαi2 knockdown leads to a decrease in microtubule dynamics, which in turn increase Rac-GTP towards the leading edge. This shift is accompanied by reduced levels of cortical actin and impaired focal adhesion disassembly, culminating in compromised cell migration. Notably, nocodazole, a microtubule-depolymerizing agent, not only diminishes Rac-GTP localization at the leading edge but also rescues cell morphology, restores normal cortical actin localization, and promotes focal adhesion disassembly, thereby facilitating cell migration. If Rac1 activity were indeed upstream of microtubules, it would be expected that nocodazole would not reduce Rac-GTP levels at the cell leading edge. These results suggest that the regulation of Rac1 activity may follow, rather than precede, alterations in microtubule dynamics, in the context of NC cells. Furthermore, in support of our model, our protein interaction analysis demonstrates Gαi2 interacting with microtubule components such as EB proteins and through this with tubulin. As we already mention above, earlier studies have reported that microtubule dynamics promote Rac1 signaling at the leading edge and by releasing RhoGEFs, promote RhoA signaling as well (Ren et al., 1999; Best et al., 1996; Garcin and Straube, 2019; Moore et al., 2013; Waterman-Storer et al., 1999). In addition, it is well-documented that RhoGEFs interact with microtubules, including βPix, a GEF for Rac1 and Cdc42, which, in turn, promotes tubulin acetylation (Kwon et al., 2020). Interestingly, in ovarian cancer cells, Gαi2 has been shown to activate Rac1 through an interaction with βPix, thereby jointly regulating migration in response to LPA (Ward et al., 2015). Taken together, these findings further support our proposed model (Fig. 6).

Therefore, in the context of collective cranial NC cells migration, our findings reveal the pivotal role played by Gαi2 in orchestrating the intricate interplay of microtubule dynamics and cellular polarity. When Gαi2 levels are diminished, we observe significant impediments in the ability of cells to efficiently navigate through their environment, resulting in a range of distinct effects. First and foremost, Gαi2 deficiency leads to the diminished ability of cells to adjust and reorient new protrusions effectively. Primary protrusions exhibit higher stability and heightened levels of active Rac1/RhoA when compared to control conditions in the leading edge. In addition, we observe a notable increase in protrusion area, a decrease in retraction velocity, and an enhanced level of cell-matrix adhesion in Gαi2 knockdown cells. These findings underscore the pivotal role that Gαi2 plays in the modulation of various cellular dynamics essential for collective cranial NC cells migration. Notably, the application of nocodazole, a microtubule-depolymerizing agent, and NSC73266, a Rac1 inhibitor, to Gαi2 knockdown cells leads to the rescue of the observed effects, thus facilitating migration. This observed response closely mirrors the outcomes associated with Par3, a known regulator of microtubule catastrophe during contact inhibition of locomotion (CIL) in NC cells (Moore et al., 2013). This parallel implies that there exists a delicate equilibrium between microtubule dynamics and Rac1-GTP levels, crucial for the establishment of proper cell polarity during collective migration. Our findings collectively position Gαi2 as a central master regulator within the intricate framework of collective cranial NC migration. This master regulator role is pivotal in orchestrating the dynamics of polarity, morphology, and cell-matrix adhesion by modulating microtubule dynamics through interactions with EB1 and EB3 proteins, described here for the first time, possible in a protein complex involving other intermediary proteins such as other microtubules accessory proteins like CLIP170, actin intermediaries, like mDia1-2, and signaling proteins such as GDIs, GAPs and GEFs, thus fostering crosstalk between the actin and tubulin cytoskeletons. This orchestration ultimately ensures the effective collective migration of cranial NC cells (Fig. 6).

## Materials and Methods

All experiments were conducted in both *Xenopus* species (*X.t* and *X.l*) using distinct concentrations of the morpholino (MO) and mRNA, as specified in each respective methodology description.

### Embryo manipulating and microinjections

*Xenopus tropicalis* and *Xenopus laevis* embryos were obtained by *in vitro* fertilization as previously described (Maldonado-Agurto et al., 2011; Toro-Tapia et al., 2018) and staged as described in Newport and Kirschner, 1982 and Nieuwkoop and Faber, 1994. Embryos were injected at the 8-cell stage as previously described (Aybar et al., 2003). MO against *Xenopus* Gαi2 was designed by GeneTools to target the 5’ UTR site of *X. tropicalis (X.t)* and *X. laevis (X.l)* transcripts (Gαi2MO: 5’-CGACACAGCCCCAGATAGTGCGT-3’). Specifically, it hybridizes with the 5’ UTR of *X. t* Gαi2 (NM_203919), 17 nucleotides upstream of the ATG start codon. For *X. l* Gαi2, the morpholino hybridizes with both isoforms described in Xenbase. It specifically targets the 5’ UTR of the Gαi2.L isoform (XM_018258962), located 17 nucleotides upstream of the ATG start codon, and the 5’ UTR of the Gαi2.S isoform (NM_001097056), situated 275 nucleotides upstream of the ATG. *X.t* and *X.l* embryos were injected with 20 and 35 ng of Gαi2MO, respectively. Equimolar concentrations of standard control MO (Control MO: 5’-CCTCTTACCTCAGTTACAATTTATA-3’) were used as previously described (Toro-Tapia et al., 2018). Fluorescein-dextran (Invitrogen, D1821; 3μg) or Rhodamine-dextran (Invitrogen, D1824; 5μg) were used as linage tracers. Plasmids were linearized and mRNA transcribed as described before (Weintraub et al., 1985). The following mRNAs were co-injected: H2B-Cherry (100 pg *X.t.*, 200 pg *X.l.*), H2B-GFP(100 pg *X.t.*, 200 pg *X.l*.), membrane-GFP (100 pg *X.t*., 200 pg *X.l*.), EB3-GFP (100 pg *X.t*., 200 pg *X.l.*), V5Gαi2 (100 pg *X.t*., 200 pg *X.l*.) and Lifeact-GFP (150 pg *X.t*., 300 pg *X.l*.). In the case of mRNA from the GTPase-based probes pGBD-GFP or rGBD-mCherry, injections were carried out the same way as described before (Leal et al., 2018).

### Whole mount *in situ* hybridization

Constructs with pSP72/snail2 and pBluescript/Twist from Labonne and Bronner-Fraser, 2000 and Mayor et al., 2000 were linearized with the restriction enzymes BglII and SmaI, respectively. For antisense RNA probe synthesis, we used Sp6 RNA Polymerase (NEB) with dig-UTP as previously described (Toro-Tapia et al., 2018). Stage 16 (for induction) or stage 23-24 (for migration) fixed embryos were processed as previously described in Maldonado-Agurto et al., 2011. Probe detection was performed with alkaline phosphatase conjugated anti-DIG antibodies (Roche) and nitroblue tetrazolium/5-bromo-4-chloro-3-indolyl phosphate (NBT/BCIP) as described before (Toro-Tapia et al., 2018). Images were collected using an Olympus SZ61 stereomicroscope and a Leica DFC450 camera from the University of Concepcion.

### Explants and microdissection

NC cells dissection was conducted following the protocol by Alfandari et al., 2003, and fibronectin dishes were prepared as previously described (Theveneau et al., 2010; Toro-Tapia et al., 2018) using 10 μg/ml of fibronectin (Sigma) for plastic dishes and 50 μg/ml for glass dishes. When required, nocodazole (Sigma, M1404) or NSC23766 (Rac inhibitor, Sigma, SML0952) were added in the DFA medium (53mM NaCl, 5mM Na_2_CO_3_, 4.5mM K-Gluconate, 32mM Na-Gluconate, 1mM MgSO_4_, 1mM CaCl_2_, 0.1% BSA and 50µg/ml streptomycin) to provide final concentrations of 10 μM-65 nM and 20 nM (20 nM for *X. laevis* and 10 nM for *X. tropicalis)*, respectively. NC explants were imaged using high magnification time-lapse on a Leica SP8 confocal microscope from University College London in UK and Centro de Microscopía Avanzada (CMA-BioBio) from University of Concepción, Chile. For low-magnification experiments, a Leica DM5500 compound microscope was utilized.

### Cell lysates and co-immunoprecipitation

Embryos from stages 23-24 were collected in lysis buffer as previously described (Toro-Tapia et al., 2018). Total protein was obtained and quantified by Bradford Method, using 25 μg for Western Blot and 0.5 mg/1 mg for co-immunoprecipitation assays. For co-immunoprecipitations, lysates were precleared with protein A/G beads (Thermo Fisher Scientific) to clean it properly. Then, they were incubated in protein A/G beads conjugated with V5 and Gαi2 antibodies, respectively. We used A/G proteins without antibody and A/G proteins with normal mouse IgG antibody as control. The immunoprecipitations were finally eluted at 37 degrees Celsius in 1% SDS for 1 hour. Proteins were separated by SDS-PAGE gels and transferred to nitrocellulose membranes. The proteins were detected using several antibodies (mentioned in the next section).

### Western blotting and immunostaining

Western blots were performed as described previously (Kuriyama and Mayor, 2008) using the following antibodies: anti Gαi2 (Santa Cruz, L-5; 1:1000); anti GAPDH (Novus Biologicals, 1D4; 1:1000); anti V5-probe (Santa Cruz, sv5-pk; 1:5000); anti acetylated tubulin (Sigma, 611B1; 1:1000); anti α-tubulin (Sigma, T9026; 1:500); anti GFP (ThermoFisher, C163; 1:500); anti EB1 (Sigma, E3406; 1:100); mouse IgG HRP (Jackson ImmunoResearch Laboratories, 1:10000); rabbit IgG HRP (Jackson ImmunoResearch Laboratories, 1:10000). Blots were revealed using a Western Lightening Plus-ECL kit (PerkinElmer) and Ultracruz Autoradiography Film Blue (Santa Cruz Biotechnology). Re-blots were performed using low pH stripping solution (1% p/v SDS, 25 mM glycine-HCl). Band intensity was measured using ImageJ (NIH). Immunostaining was performed previously described (Toro-Tapia et al., 2018) using the following antibodies: α-tubulin (Sigma, T9026; 1:100), acetylated tubulin (Sigma, 611B1; 1:100), Par3 (Santa Cruz, sc-5598; 1:20), ζ-PKC (Santa Cruz, (B-7), sc-393218; 1:20), anti-phosphopaxilin (Invitrogen, pY118; 1:20), anti β-integrin (DSHB, 8C8; 1:10), anti N-cadherin (DSHB, 5D3; 1:10), rabbit IgG Alexa 488 (Invitrogen, A11034; 1:100) and mouse IgG Alexa 488 (Invitrogen, A11017; 1:200) antibodies. Hoescht (Thermo Fischer Scientific, 1:1000), Phalloidin 546 and Phalloidin 633 (Life Technologies, 1:500) were applied with the secondary antibody. The imaging was performed in a Zeiss LSM780 Confocal Microscope using 40X and 63X oil objectives (CMA-BioBio), University of Concepción.

### *In situ* proximity ligation assay (PLA)

For the *in situ* visualization of Gαi2-protein interactions, we followed the Duolink PLA Fluorescence Protocol from Sigma-Aldrich, using the Duolink *In Situ* Detection Reagents Red kit (DUO92008, Sigma). Briefly, NC cells were cultured on fibronectin-coated coverslips and fixed in 3.7% formaldehyde in PBS for 30 minutes. Permeabilization was performed with 0.3% Triton-X in PBS for 15 minutes, followed by PBS washes. NC cells were then blocked in a Blocking Solution (Sigma-Aldrich) at 37 °C for one hour and incubated with primary antibodies: α-tubulin (Sigma, T9026; 1:200), Gαi2 (Santa Cruz, L-5, and T-19; 1:50), EB1 (Sigma, E3406; 1:100), all diluted in Duolink Antibody Diluent (Sigma-Aldrich).

After washing in wash Buffer A (10 mM Tris, 150 mM NaCl, and 0.05% Tween 20), NC cells were incubated with secondary antibodies conjugated with PLUS and MINUS probes (Sigma-Aldrich) for one hour at 37 °C. The following Duolink *in Situ* PLA secondary antibodies were used: Anti-Mouse MINUS (DUO92004; 1:5, Sigma-Aldrich), Anti-Rabbit PLUS (DUO92002; 1:5, Sigma-Aldrich).

Subsequently, NC cells were washed three times in Buffer A and incubated in 1 unit/μl of ligase in diluted Ligase Buffer (1:5, Sigma Aldrich) for 30 minutes at 37 °C. After three washes in Buffer A, cells were incubated in 5 units/μl of DNA polymerase in diluted Polymerase Buffer with Red Fluorescent-labeled Oligonucleotides (1:5, Sigma Aldrich) for 100 minutes at 37 °C. A final round of washes was performed twice for 10 minutes in 1X Buffer B, then 1 minute in 0.01X Buffer B. The slides were then mounted using a 5 μl volume of Duolink *In Situ* mounting medium containing DAPI. Nail polish was used to seal the edges of the coverslip. Negative controls were performed using secondary antibodies only. Imaging was carried out on a Leica SP8 confocal microscope using a 63X oil objective (CMA-BioBio), University of Concepción.

### Cell dispersion and morphology

Cell dispersion was studied by time-lapse using DMX6000 Zeiss microscope from University of Concepción equipped with a motorized stage and a Leica camera (DFL420) with a N Plan ApoChromat 20x/0.25 NA objective controlled by LAS EZ software. Frames were obtained every 3 minutes for 4 hours in *X.t* and every 5 minutes for 10 hours in *X.l*. The dispersion each initial size of the explant was normalized upon the control and rate was quantified by Delaunay triangulation using a plugin for ImageJ as previously described (Carmona-Fontaine et al., 2011), comparing the final and initial average area for each condition. Every Delaunay triangulation calculates the area generated between three adjacent cells and it grows depending on how much disperse are the cells between each other in the final point. For cell morphology time-lapse, the images were acquired using a 40x objective in a SP8 Confocal Microscope (University College London), tracking cells injected with membrane-GFP and lifeactin-GFP. For actin dynamics, the images were acquired using a 63X objective in an inverted SP8 Confocal Microscope (CMA-BioBio from University of Concepción). The frames were collected every 20s for 30 minutes in total and processed subtracting frame per frame to measure extension/retraction area and using ADAPT plugin from ImageJ to calculate extension/retraction perimeter and extension/retraction speed (Barry et al., 2015).

### Microtubule dynamics analysis

EB3-GFP images were acquired at intervals of 1.5 or 3 seconds per frame for a total duration of 5 minutes, utilizing the SP8 Confocal Microscope from CMA-BioBio at the University of Concepción. Temperature conditions were regulated using an incubator chamber, set to 28 degrees Celsius for *X.t*. and 23 degrees Celsius for *X.l*. The recorded movies were analyzed automatically through plusTipTracker (Applegate et al., 2011), as previously described (Moore et al., 2013). The PlusTipTracker software separates comets based on their speed and lifetime, classifying them as fast long-lived, fast short-lived, slow long-lived, or slow short-lived. In both conditions (control and morphant cells), we compared the percentage of each type of comet, as previously described in Moore et al., 2013. The following parameters were applied: 8 frames as the maximum gap length, 3 frames as the minimum track length, 5-25 pixels as the search radius range, 35 as the maximum forward angle, 10 as the maximum backward angle, 1.5 as the maximum shrinkage factor and a 2-pixel fluctuation radius. For MtrackJ analysis, 5×5 µm squares were selected at the front and back of a cell. The total number of EB3-GFP +tips that disappeared within the square during a 2 minute period was compared with the number of EB3-GFP +tips entered to the square. Microtubules present within the square in the final frame were excluded from the analysis.

### Statistical analysis

The normality of data sets was assessed using Kolmogorov-Smirnov’s test, D’Agostino-Pearson test, and Shapiro-Wilks tests in Prism6 (GraphPad). For data sets that exhibited a normal distribution, we employed the T-student test (two-tail) or one-way ANOVA for comparison. In cases where the data sets did not follow a normal distribution, such as in cell dispersion assays, we utilized Mann-Whitney’s test or nonparametric ANOVA (Kruskal-Wallis) in Prism6. Cross-comparisons were conducted only if the overall P-value of the ANOVA was less than 0.1.

## Acknowledgements

This work was supported by grants from the Fondo Nacional de Desarrollo Cientifico y Tecnologico (FONDECYT) 1180926 (MT), Vicerrectoria de Investigacion UdeC (VRID) 2022000483INV (MT), Faculty of Biological Sciences UdeC FCB-I-2022-03 (MT) and from the Medical Research Council (MR/S007792/1) (RM), Biotechnology and Biological Sciences Research Council (M008517, BB/T013044) (RM) and Wellcome Trust (102489/Z/13/Z) (RM). We also acknowledge the Agencia Nacional de Investigacion y Desarrollo (ANID) and Direccion de postgrado UdeC for supporting the graduate studies of SV, JIL, LMT, MJR and JG. We acknowledge Centro de Microscopia Avanzada (CMA-Bio-Bio) for supporting with imaging acquisition and finally, we thank Gustavo Minder for the artwork.

## Supplementary material

**Supplementary Movie S1. Gαi2 is required for cranial NC cell migration *in vitro* in *Xenopus*.**

**Supplementary Movie S2. 3D reconstruction of microtubules cytoskeleton in control and Gαi2 knockdown conditions.**

**Supplementary Movie S3. Gαi2 morpholino disrupt microtubule dynamics.**

**Supplementary Movie S4. Gαi2 knockdown condition affects active Rac1 localization during CIL in cranial NC cells.**

**Supplementary Movie S5. Rac1 inhibitor partially restore cranial NC cell migration in Gαi2 knockdown conditions.**

**Supplementary Movie S6. Gαi2 morphant cells show an increase in the focal adhesion stability.**

**Supplementary Movie S7. Nocodazole restore *in vitro* cranial NC cell migration in Gαi2 knockdown conditions.**

**Supplementary Movie S8. Gαi2MO affects cell morphology and nocodazole can restore cell morphology under morphant conditions.**

---

**Supplementary Figure S1.**
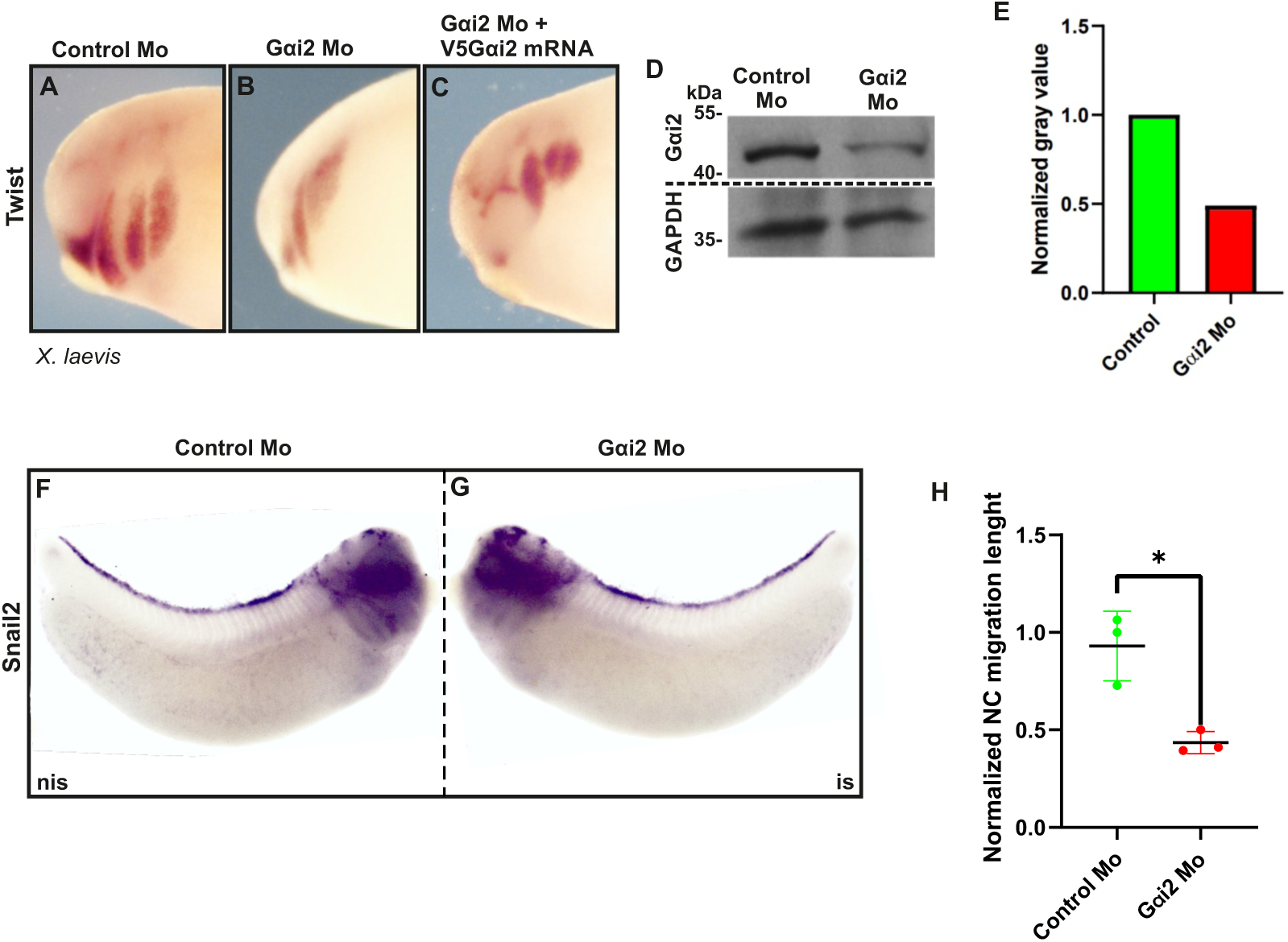
Gαi2 knockdown affect *Xenopus laevis* migration *in vivo*. **A-C.** *In situ* hybridization against *twist* performed in *X.l*. embryos at stage 25-26 to analyze *in vivo* early migration in control, morphant and recued embryos. N= 18 embryos per condition. **D-E.** Western blot of lysates from *X.l.* embryos injected with Gαi2 morpholino shows efficiently Gαi2 knockdown. **F-G.** *In situ* hybridization against *snail2* performed in *X.l.* embryos at stage 32 to analyze late *in vivo* migration in controls and morphant embryos. N= 3 embryos per condition. **I. H.** Quantification of the migration length in **F** and **G**. Each point corresponds to one embryo analyzed. Error bars are ± s.e.m. P*≤0.05 (two-tailed Student’s t-test).

**Supplementary Figure S2.**
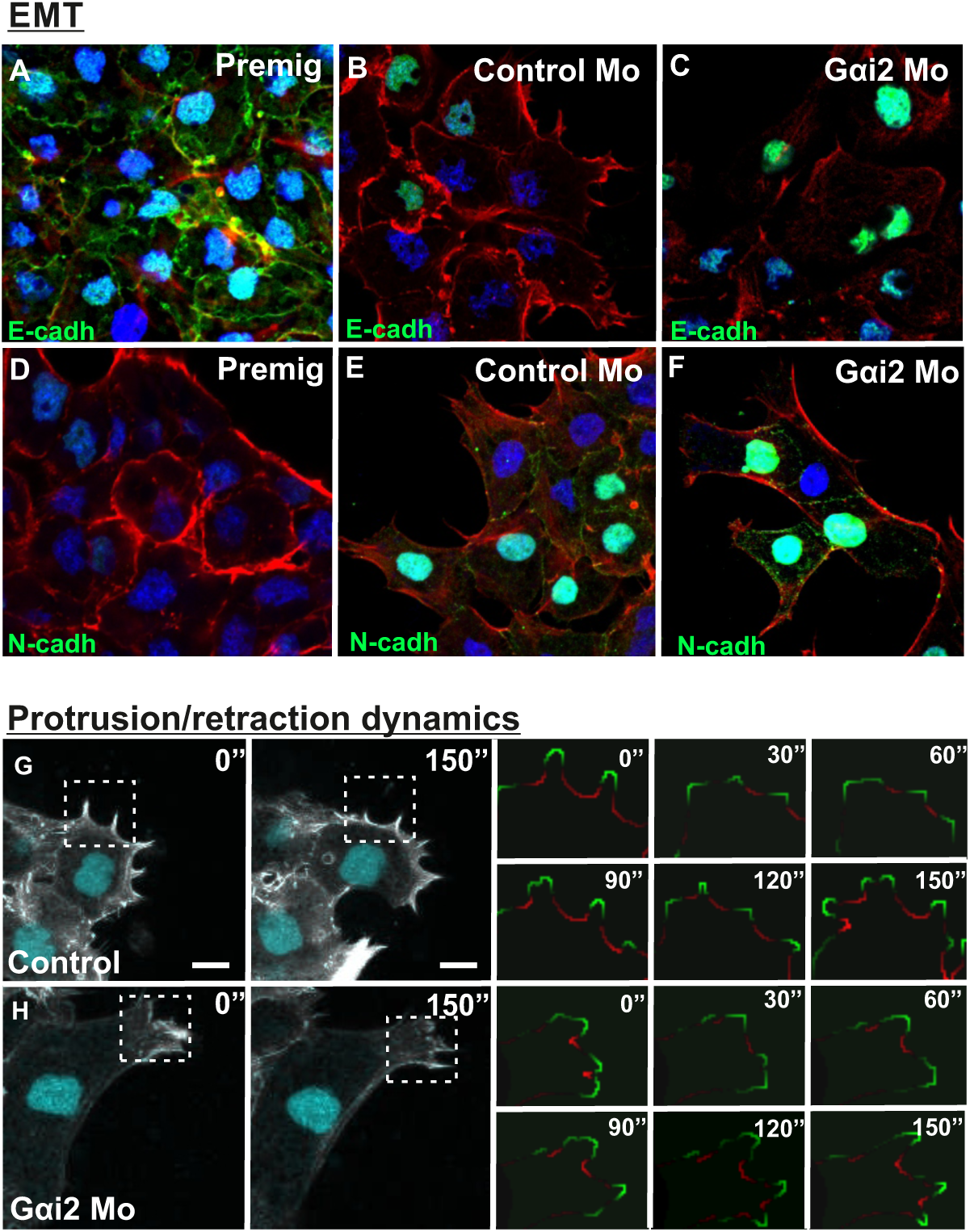
Gαi2MO does not affect cadherin switching during EMT. **A-F.** Immunofluorescence assays were performed in cranial NC cell explants from *X.t* using anti-E-cadherin (E-cadh: green, A-C) and anti-N-cadherin (N-cadh: green, D-F) to detect the cadherin switching. E-cadherin is localized at cell-cell contact on pre-migratory cell explants (A) and N-cadherin is localized at cell-cell contact on migratory explants from Control MO (E) and Gαi2 knockdown conditions (F). Phalloidin is showed in red and nuclei were stained with Hoechst (blue). **G-H.** Example of protrusion and retraction dynamics measurements using ADAPT plugin for ImageJ. The area of interest was segmented. Protrusion dynamics are showed in green and retraction dynamics are showed in red. Scale bar: 10 μm.

**Supplementary Figure S3.**
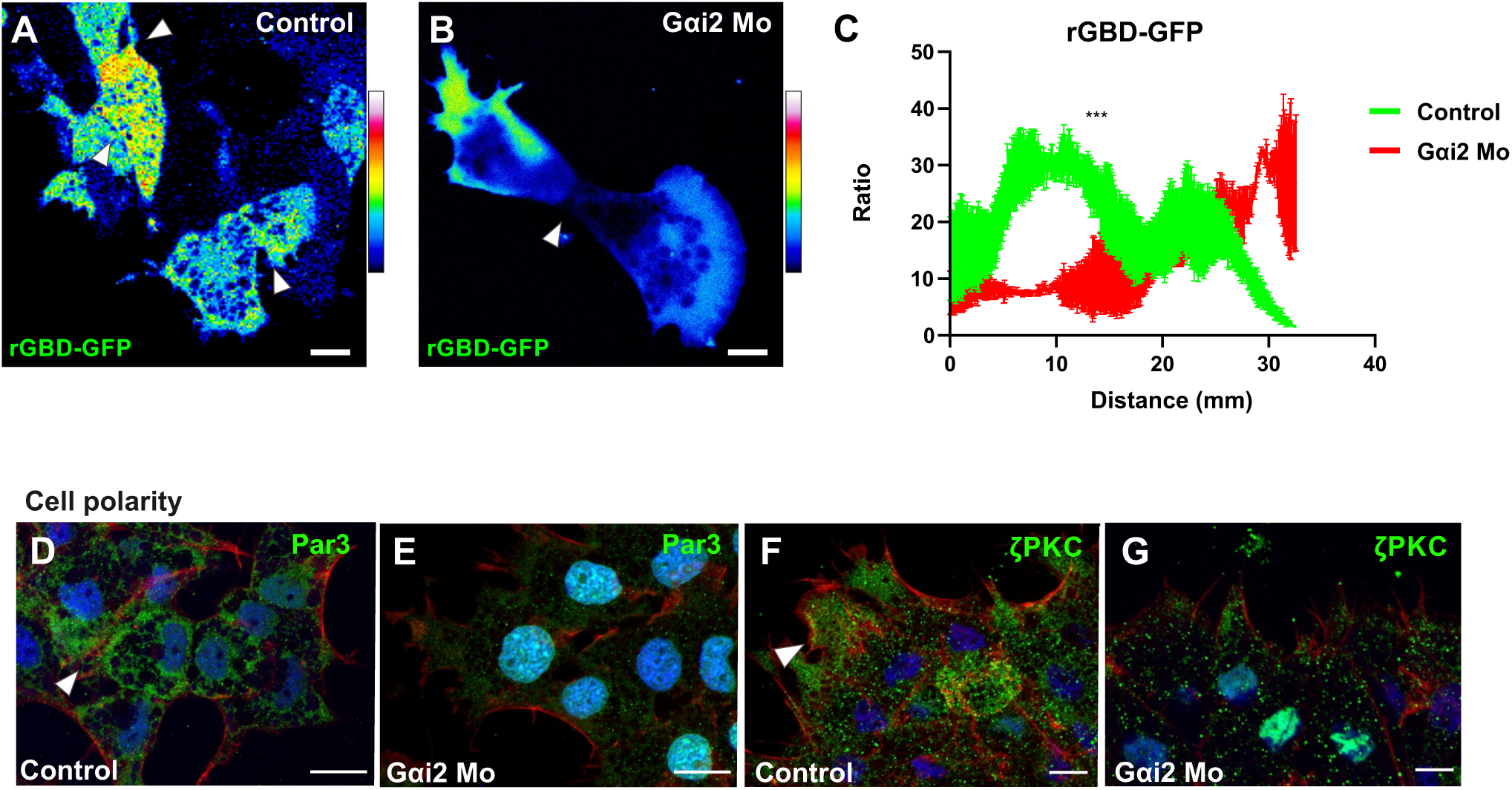
Gαi2 knockdown affect active RhoA-GTP, and the polarity markers Par3 and ζ-PKC localization. **A-B.** Pseudocolor scale for visualization of changes in the active RhoA localization using rGBD-GFP probe. This probe contains the RhoA binding domain of the effector protein Rhotekin fused to GFP, which makes it possible to observe the localization of active RhoA over time. In control cells active RhoA is highly concentrated in the cell-cell contact. In the Gαi2 morphant cells active RhoA disappear in the cell-cell contact, increasing its localization in the cell protrusion. **C.** Quantification of active RhoA fluorescence intensity in time, showing a significant contraposition in RhoA localization between control and morphant condition. The significance was evaluated with a Mann-Whitney test for non-parametric data (***, P<0.001). Error bars: s.e.m. Magnification: 60X. (experimental N = 5). Scale bar: 10 μm. **D-E.** Immunofluorescence against Par3 (green). Red: phalloidin, blue: Hoechst. Gαi2MO delocalizes Par3 from cell-cell contact. Scale bar: 10 μm. **F-G.** Immunofluorescence against ζ-PKC (green). Red: phalloidin, blue: Hoechst. Gαi2MO delocalizes ζ-PKC from cell cortex.

**Supplementary Figure S4.**
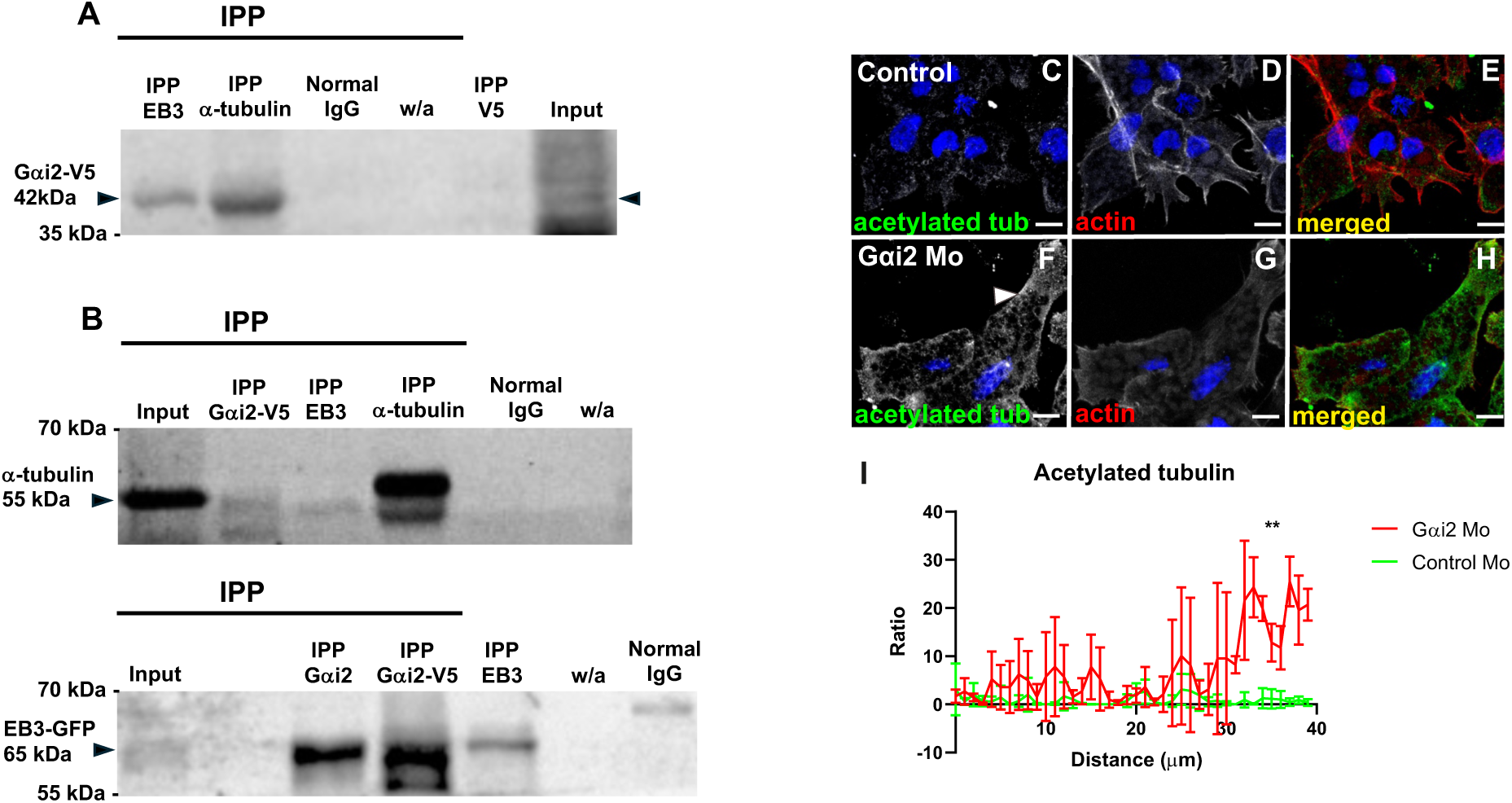
Gαi2 conform a microtubules interaction complex in cranial NC cells and increases acetylated tubulin signal in the protrusions. **A-B.** Duplicate of coimmunoprecipitations showing the interaction between Gαi2-V5, α-tubulin and EB3. IPP EB3: immunoprecipitation conjugated to GFP antibody (EB3-GFP). IPP α-tubulin: immunoprecipitation conjugated to α-tubulin antibody. IPP Gαi2-V5: immunoprecipitation conjugated to V5 antibody (Gαi2-V5). Normal IgG: non-related IgG. w/a: Without antibody. All embryos were injected with Gαi2-V5. IPP V5: immunoprecipitation conjugated to V5 antibody alone, from embryos injected with V5, as a control **(A)**. Western blot performed against V5 **(A)** and a-tubulin and GFP antibodies **(B)**. n total = 2 lysates per condition. **C-H.** Immunofluorescence against acetylated tubulin (green) and actin (red) in control and Gαi2 morphant explants to localize stable microtubules in *X.t*. Under Gαi2 knockdown conditions, acetylated tubulin increases towards the leading edge. Scale bar: 10 μm. **I.** Quantification of acetylated tubulin fluorescence intensity from nucleus to leading edge, showing a strong concentration of acetylated tubulin towards the leading edge. The significance was evaluated with a Mann-Whitney test for non-parametric data (**, P<0.01).

## References

Abercrombie M, Heaysman JE, Pegrum SM (1971) The locomotion of fibroblasts in culture IV Electron microscopy of the leading lamella. Exp Cell Res 67: 359–367

Akhmanova A and Steinmetz MO (2015) Control of microtubule organization and dynamics: two ends in the limelight. Nature Reviews Molecular Cell Biology 12: 711–726

Akhshi TK, Wernike D, Piekny A (2013) Microtubules and actin crosstalk in cell migration and division. Cytoskeleton. 71: 1–23

Alfandari D, Cousin H, Gaultier A, Hoffstrom BG, DeSimone DW. (2003). Integrin α5B1 supports the migration of *Xenopus* cranial neural crest in fibronectin. Dev Biol. 260: 449–464

Applegate KT, Besson S, Matov A, Bagonis MH, Jaqaman K and Danuser G (2011) PlusTipTracker: Quantitative image analysis software for the measurement of microtubule dynamics. Journal of Structural Biology 176: 168–184

Aybar MJ, Nieto MA, Mayor R. (2003). Snail precedes Slug in the genetic cascade required for the specification and migration of the *Xenopus* neural crest. Development 130: 483–494

Barriga EH, Franze K, Charras G, Mayor R. (2018) Tissue stiffening coordinates morphogenesis by triggering collective cell migration *in vivo*. Nature 554: 523–527

Barry DJ, Durkin CH, Abella JV, Way M (2015) Open source software for quantification of cell migration, protrusions, and fluorescence intensities. J Cell Biol. 209:163–80

Bershadsky A., Chausovsky A., Becker E., Lyubimova A., Geiger B (1996) Involvement of microtubules in the control of adhesion-dependent signal transduction. Curr Biol. 6: 1279–89.

Best A, Ahmed S, Kozma R and Lim L (1996) The Ras-related GTPase Rac1 binds tubulin. J. Biol. Chem 271: 3756 –3762

Bieling P, Laan L, Schek H, Munteanu EL, Sandblad L, Dogterom M, Brunner D and Surrey T (2007) Reconstitution of a microtubule plus-end tracking system in vitro. Nature 450: 1100–1105

Bouchet BP and Akhmanova A (2017) Microtubules in 3D cell motility. J. Cell Sci. 130: 39–50

Bouchet BP, Gough RE, Ammon YC, van de Willige D, Post H, Jacquemet G, Altelaar AFM, Heck AJR, Goult BT, Akhmanova A (2016) Talin-KANK1 interaction controls the recruitment of cortical microtubule stabilizing complexes to focal adhesions. eLife 5, e18124

Bronner-Fraser M, Sauka-Spengler T (2008) Evolution of the neural crest view from a gene regulatory perspective. Genesis 46: 673–682

Carmona-Fontaine C, Theveneau E, Tzekou A, Tada M, Woods M, Page KM, Parsons M, Lambris JD and Mayor R (2011) Complement fragment C3a controls mutual cell attraction during collective cell migration. Developmental Cell 21: 1026–1037

Chang WK, Carmona-Fontaine C and Xavier JB (2013) Tumor–stromal interactions generate emergent persistence in collective cancer cell migration. Interface Focus 3: 20130017–20130017

Chao WT and Kunz J (2009) Focal adhesion disassembly requires clathrin-dependent endocytosis of integrins. FEBS Lett. 583: 1337–1343

Chen X and Macara IG (2005) Par-3 controls tight junction assembly through the Rac exchange factor Tiam1. Nature Cell Biology 7: 262–269

Cowan CR and Hyman AA (2007) Acto-myosin reorganization and PAR polarity in C. elegans. Development 134: 1035–1043

Cotton M and Claing A (2009) G protein-coupled receptors stimulation and the control of cell migration. Cell. Signal. 21:1045–1053

Dogterom M and Koenderink GH (2019) Actin–microtubule crosstalk in cell biology Nature Reviews Molecular Cell Biology 1: 38–54.

Dubois F, Alpha K, Turner CE (2017) Paxillin regulates cell polarization and anterograde vesicle trafficking during cell migration. Mol Biol Cell. 28: 3815–3831

Eshun-Wilson L, Zhangb R, Portran D, Nachury MV, Toso DB, Löhr T, Vendruscolo M, Bonomi M, Fraser JS and Nogales E (2019) Effects of α-tubulin acetylation on microtubule structure and stability. Proc Natl Acad Sci U S A 116: 10366–10371

Etienne-Manneville S (2013) Microtubules in cell migration. Annu. Rev. Cell Dev. Biol. 29: 471–499

Ezratty EJ, Partridge MA and Gundersen GG (2005) Microtubule-induced focal adhesion disassembly is mediated by dynamin and focal adhesion kinase Nat. Cell Biol. 7: 581–590

Fuentealba J, Toro-Tapia G, Arriagada C, Riquelme L, Beyer A, Henriquez JP, Caprile T, Mayor R, Marcellini S, Hinrichs MV, Olate J and Torrejon M (2013) Ric-8A, a guanine nucleotide exchange factor for heterotrimeric G proteins, is critical for cranial neural crest cell migration. Dev. Biol. 378, 74e82

Fuentealba J, Toro-Tapia G, Rodriguez M, Arriagada C, Maureira A, Beyer A, Villaseca S, Leal JI, Hinrichs MV, Olate J, Caprile T, Torrejón M (2016) Expression profiles of Gαsubunits during *Xenopus tropicalis* embryonic development. Gene Expression Pattern 22: 15–25

Galjart N (2010) Plus-end-tracking proteins and their interactions at microtubule ends. Current Biology 20: 528–537

Garcin C and Straube A (2019) Microtubules in cell migration. Essays Biochem. 63: 509–520

Gotta M and Ahringer J (2001) Distinct roles for Galpha and Gbetagamma in regulating spindle position and orientation in Caenorhabditis elegans embryos. Nat. Cell Biol. 3: 297–300

Gotta M, Abraham MC and Ahringer J (2001) CDC-42 controls early cell polarity and spindle orientation in C. elegans. Curr. Biol. 11: 482–488

Gu Z, Noss EH, Hsu VW, Brenner MB. (2011) Integrins traffic rapidly via circular dorsal ruffles and micropinocytosis during stimulated cell migration. J Cell Biol. 193:61–70

Han SB, Moratz C, Huang NN, Kelsall B, Cho H, Shi CS et al (2005) Rgs1 and Gnai2 regulate the entrance of B lymphocytes into lymph nodes and B cell motility within lymph node follicles. Immunity 22: 343–354

Hao Y, Du Q, Chen X, Zheng Z, Balsbaugh JL, Maitra S, Shabanowitz J, Hunt DF, Macara IG (2010) Par3 controls epithelial spindle orientation by aPKC-mediated phosphorylation of apical Pins. Curr Biol 20: 1809–1818

Heath JP and Dunn GA (1978) Cell to substratum contacts of chick fibroblasts and their relation to the microfilament system. A correlated interference-reflexion and highvoltage electron-microscope study. J Cell Sci 29: 197–212

Henty-Ridilla JL, Rankova A, Eskin JA, Kenny K, Bruce L and Goode BL (2016) Accelerated actin filament polymerization from microtubule plus ends. Science. 352: 1004–1009

Huttenlocher A, Sandborg RS and Horwitz AF (1995) Adhesion in cell migration. Curr Opin Cell Biol 7: 697– 706

Hwang IY, Park C and Kehrl JH (2007) Impaired trafficking of Gnai2+/– and Gnai2–/– T lymphocytes: implications for T cell movement within lymph nodes. J. Immunol. 179: 439–448

Hwang IY, Park C, Harrison K, Boularan C, Galés C and Kehrl JH (2015) An Essential Role for RGS Protein/Gα_i2_ Interactions in B Lymphocyte–Directed Cell Migration and Trafficking. J Immunol. 194: 2128–2139

Ju RJ, Falconer AD, Dean KM, Fiolka RP, Sester DP, Nobis M, Timpson P, Lomakin AL, Danuser G, White MD, Oelz DB, Haass NK and Stehbens SJ (2022) Compression-dependent microtubule reinforcement comprises a mechanostat which enables cells to navigate confined environments. BioRxiv. 10.1101/2022.02.08.479516

Kaverina I, Krylyshkina O and Small JV (1999) Microtubule targeting of substrate contacts promotes their relaxation and dissociation. Journal of Cell Biology 146: 1033–1043

Kelleher FC, Fennelly D and Rafferty M (2006) Common critical pathways in embryogenesis and cancer. Acta. Oncol. 45: 375–388

Kiyomitsu T (2019) The cortical force-generating machinery: how cortical spindlepulling forces are generated. Curr. Opin. Cell Biol. 60: 1–8

Kjoller L and Hall A (1999) Signaling to Rho GTPases *Exp*. Cell Res. 253: 166–179

Krendel M, Zenke FT, Bokoch GM. (2002). Nucleotide exchange factor GEF-H1 mediates cross-talk between microtubules and the actin cytoskeleton. Nat Cell Bio. 4: 294–301

Kuriyama S and Mayor M (2008) Molecular analysis of neural crest migration. Phil. Trans R Doc. 363: 1349–1362

Laan L, Husson J, Munteanu EL, Kerssemakers JW and Dogterom M (2008) Force-generation and dynamic instability of microtubule bundles. Proc. Natl. Acad. Sci. USA 105: 8920–8925

LaBonne C, Bronner-Fraser M. (2000). Snail-related transcriptional repressors are required in *Xenopus* for both the induction of the neural crest and its subsequent migration. Dev Biol. 221: 195–205

Kwon Y, Jeon YW, Kwon M, Cho Y, Park D, Shin JE. (2020). βPix-d promotes tubulin acetylation and neurite outgrowth through a PAK/Stathmin1 signaling pathway. PloS ONE. 15: e0230814

Lauffenburger DA and Horwitz AF (1996) Cell migration: A physically integrated molecular process. Cell 84: 359–369

Leal JI, Villaseca S, Beyer A, Toro-Tapia G and Torrejón M (2018) Ric-8A, a GEF for heterotrimeric G-proteins, controls cranial neural crest cell polarity during migration. Mechanisms of Development 154: 170–178

Lee CC, Cheng YC, Chang CY, Chi-Min Lin CM and Chang JY (2018) Alpha-tubulin acetyltransferase/MEC-17 regulates cancer cell migration and invasion through epithelial–mesenchymal transition suppression and cell polarity disruption. Sci Rep. 8, 17477

Li H, Yang L, Fu H, Yan J, Wnag Y and Guo H (2013) Association between Gαi2 and ELMO1/Dock180 connects chemokine signalling with Rac activation and metastasis. Nat. Commun. 4, 1706

Li Z, Cai S, Liu Y, Yang CL, Tian Y, Chen G et al (2016) Over-expression of Gαi3 in human glioma is required for Akt-mTOR activation and cell growth. Oncotarget 5, 1

Liu Y, Seto E, Kuo E, Yu W, Schartz RJ, Blazo M, Zhang SL, Peng X (2013) Inactivation of Cdc42 in neural crest cells causes craniofacial and casdiovascular morphogenesis defects. Dev. Biol.

Liu Y, Chen MB, Cheng L, Zhang ZG, Yu ZQ, Jiang Q et al (2018) microRNA-200a downregulation in human glioma leads to Gαi1 over-expression, Akt activation, and cell proliferation. Oncogene 37: 2890–2902

Marchant CL, Malmi-Kakkada AN, Espina JA and Barriga EH (2022) Cell clusters softening triggers collective cell migration in vivo. Nat Mater. 21: 1314–1323

Marchant CL, Malmi-Kakkada AN, Espina JA and Barriga EH (2021) Microtubule deacetylation reduces cell stiffness to allow the onset of collective cell migration in vivo. BioRxiv 10.1101/2021.08.12.456059

Matthews HK, Marchant L, Carmona-Fontaine C, Kuriyama S, Larrain J, Holt AR, Parsons M and Mayor R (2008) Directional migration of neural crest cells in vivo is regulated by Syndecan-4/Rac1 and non-canonical Wnt signaling/RhoA. Development 135: 1771–1780

Mayor R and Etienne-Manneville S (2016) The front and rear of collective cell migration. Nature Review of Molecular Cell Biology 17: 97–109

Mayor R, Guerrero N, Young RM, Gomez-Skarmeta JL, Cuellar C. (2000). A novel function for the Xslug gene: control of dorsal mesoderm development by repressing BMP-4. Mech of Dev. 97: 47–56

Mayor R and Theveneau E (2013) The neural crest. Development 140: 2247–2251

Mimori-Kiyosue Y, Shiina N and Tsukita S (2000) The dynamic behavior of the APC-binding protein EB1 on the distal ends of microtubules. Current biology 14: 865–868

Mitra SK, Hanson DA and Schlaepfer DD (2005) Focal adhesion kinase: in command and control of cell motility. Nat Rev Mol Cell Biol. 6: 56–68

Molina NA, Rodrigues-Ferreira S, Honoré S and Nahmias C (2017) Regulation of end-binding protein EB1 in the control of microtubule dynamics. Cellular and Molecular Life Sciences 13: 2381–2393

Moore R, Theveneau E, Pozzi S, Alexandre P, Richardson J, Merks A et al (2013) Par3 controls neural crest migration by promoting microtubule catastrophe during contact inhibition of locomotion. Development 140: 4763–4775

Newport J, Kirschner M. (1982). A major developmental transition in early Xenopus embryos: II. Control of the onset of transcription. Cell. 30: 687–96

Nieuwkoop PD, Faber J. (1967). Normal table of Xenopus laevis (Doudin). Amsterdam: Elsevier-North Holland Publishing

Nobes CD and Hall A (1995) Rho, Rac, and Cdc42 GTPases regulate the assembly of multimolecular focal complexes associated with actin stress fibers, lamellipodia, and filopodia Cell 81: 53–62

Nůsková H, Serebryakova MV, Ferrer-Caelles A, Sachsenheimer T, Luchtenborg C, Miller AK et al (2021) Stearic acid blunts growth-factor signaling via oleoylation of GNAI proteins. Nat Comm. 12, 4590

Orr AW, Pallero MA and Murphy-Ullrich JE (2002) Thrombospondin stimulates focal adhesion disassembly through Gi- and phosphoinositide 3-kinase-dependent activation. J. Biol. Chem. 277: 20453–20460

Orr AW, Pedraza CE, Pallero MA, Elzie CA, Goicoechea S, Strickland DK and Murphy-Ullrich JE (2003) Low density lipoprotein receptor–related protein is a calreticulin coreceptor that signals focal adhesion disassembly. J Cell Biol. 161: 1179–1189

Parmentier ML, Woods D, Greig S, Phan PG, Radovic A, Bryant P et al (2000) Rapsynoid/partner of inscuteable controls asymmetric division of larval neuroblasts in Drosophila. J. Neurosci. 20: 84

Parsons SA, Sharma R, Roccamatisi DL, Zhang H, Petri B, Kubes P, Colarousso P, Patel KD. (2012). Endothelial paxillin and focal adhesion kinase (FAK) play a critical role in meutrophil transmigration. Eur J Immunol. 42: 436–446

Pegtel DM, Ellenbroek S, Mertens A, van der Kammen RA, de Rooij J and Collard JG (2007) The Par-Tiam1 complex controls persistent migration by stabilizing microtubule-dependent front-rear polarity. Curr. Biol. 17: 1623–1634

Perez F, Diamantopoulos GS, Stalder R and Kreis TE (1999) CLIP-170 highlights growing microtubule ends in vivo. Cell 96: 517–527

Pero RS, Borchers MT, Spicher K, Ochkur SI, Sikora L, Rao SP et al (2007) Galphai2-mediated signaling events in the endothelium are involved in controlling leukocyte extravasation. Proc. Natl. Acad. Sci. USA 104: 4371–4376

Piacentino ML, Li Y and Bronner ME (2020) Epithelial-to-mesenchymal transition and different migration strategies as viewed from the neural crest. Curr Opin Cell Biol. 66: 43–50

Ren XD, Kiosses WB, Schwartz MA. (1999). Regulation of the small GTP-binding protein Rho by cell adhesion and the cytoskeleton. EMBO J. 18: 578–585

Ridley AJ (2011) Life at the leading edge. Cell 145: 1012–1022

Ridley AJ (2015) Rho GTPase signaling in cell migration. Curr Op Cell Biol. 36: 103–112

Ridley AJ, Schwartz MA, Burridge K, Firtel RA, Ginsberg MH, Borisy G, Horwitz AR (2003) Cell migration: Integrating signals from front to back. Science 302: 1704–1709

Rodriguez OC, Schaefer AW, Mandato CA, Forscher P, Bement WM and Waterman-Storer CM (2003) Conserved microtubule-actin interactions in cell movement and morphogenesis. Nat. Cell Biol. 5: 599–609

Rohde LA and Heisenberg CP (2007) Zebrafish gastrulation: cell movements, signals, and mechanisms. Int. Rev. Cytol. 261: 159 – 192

Rottner K and Schaks M (2019) Assembling actin filaments for protrusion. Current Opinion in Cell Biology 56: 53–63

Roychowdhury S, Panda D, Wilson L and Rasenick MM (1999) G protein alpha subunits activate tubulin GTPase and modulate microtubule polymerization dynamics. J Biol Chem. 274: 13485–13490

Roychowdhury S and Rasenick MM (2008) Submembraneous microtubule cytoskeleton: regulation of microtubule assembly by heterotrimeric Gproteins FEBS J 275:4654–63

Sah VP, Seasholtz TM, Sagi SA and Brown JH (2000) The role of Rho in G protein-coupled receptor signal transduction. Annu. Rev. Pharmacol. Toxicol. 40: 459–489

Schaefer M, Petronczki M, Dorner D, Forte M, and Knoblich, JA (2001) Heterotrimeric G proteins direct two modes of asymmetric cell division in the Drosophila nervous system. Cell 107: 183–194

Schaks M, Giannone G and Rottner K (2019) Actin dynamics in cell migration. Essays Biochem. 63: 483– 495

Seetharaman S and Etienne-Manneville S (2020) Cytoskeletal Crosstalk in Cell Migration. Trends in Cell Biology. 30: 720–735

Serre L, Stoppin-Mellet V and Arnal I (2019) Adenomatous polyposis coli as a scaffold for microtubule end-binding proteins. Journal of molecular biology 431:10

Shoval I and Kalcheim C (2012) Antagonistic activities of Rho and Rac GTPases underlie the transition from neural crest delamination to migration. Dev. Dyn. 241: 1155–1168

Steventon B and Mayor R (2012) Early neural crest induction requires and initial inhibition of Wnt signals. Dev. Biol. 367:55–65

Szabó A and Mayor R (2018) Mechanisms of neural crest migration. Ann. Rev. Gen. 52: 43–63

Theveneau E and Mayor R (2012) Neural crest delamination and migration: from epithelium-to-mesenchyme transition to collective cell migration. Dev. Biol. 366: 34– 54

Theveneau E, Marchant L, Kuriyama S, Gull M, Moepps B, Parsons M and Mayor R (2010) Collective chemotaxis requires contact-dependent cell polarity. Developmental Cell 19: 39–53

Thompson BD, Jin Y, Wu KH, Colvin RA, Luster AD and Birnbaumer L (2007) Inhibition of G alpha i2 activation by G alpha i3 in CXCR3-mediated signaling. J. Biol. Chem. 282: 9547–9555

Toro-Tapia G, Villaseca S, Beyer A, Roycroft A, Marcellini S, Mayor R and Torrejón M (2018) The Ric-8A/Gα13/FAK signaling cascade controls focal adhesion formation during neural crest cell migration. Development. 145: 1–12

Trainor PA (2010) Craniofacial Birth Defects: The Role of Neural Crest Cells in the Etiology and Pathogenesis of Treacher Collins Syndrome and the Potential for Prevention. Am J Med Genet. 12: 2984– 2994

Tucker RP, Hagios C, Chiquet-Ehrismann R, Lawler JL, Hall RJ and Erickson C (1999) Thrombospondin-1 and Neural Crest Cell Migration. Dev. Dyn. 214: 312–322

Vedula SR, Ravasio A, Lim CT and Ladoux B (2013) Collective cell migration: A mechanistic perspective. Physiology 28: 370–379

Villaseca S, Romero G, Ruiz MJ, Pérez C, Leal JI, Tovar LM and Torrejón M (2022) Gαi protein subunit: A step toward understanding its non-canonical mechanisms. Front Cell Dev Biol. 10: 941870

Wang H, Ng KH, Qian H, Siderovski DP, Chia W, Yu F. (2005). Ric-8 controls Drosophila neural progenitor asymmetric division by regulating heterotrimeric G proteins. Nat. Cell Biol. 7: 1091–1098. doi:10. 1038/ncb1317

Ward JD, Ha JH, Jayaraman M and Dhanasekaran DN (2015) LPA-mediated migration of ovarian cancer cells involves translocalization of Gαi2 to invadopodia and association with Src and β-pix. Cancer Letters 356: 382–391

Watanabe T, Noritake J, Kaibunchi K. (2005). Regulation of microtubules in cell migration. Trends Cell Biol. 15: 76–83

Waterman-Storer CM, Worthylake RA, Liu BP, Burridge K and Salmon ED (1999) Microtubule growth activates Rac1 to promote lamellipodial protrusion in fibroblasts. Nat Cell Biol 1:45–50

Weintraub H, Izant JG, Harland RM. (1985). Antisense RNA as a Molecular Toll for genetic analysis. Trends in Genetics. 1: 23–25.

Wiege K, Le DD, Syed SN, Ali SR, Novakovic A, Beer-Hammer S, Piekorz RP, Schmidt RE, Nürnberg B and Gessner JE (2012) Defective macrophage migration in Gαi2-but not Gαi3-deficient mice. J Immunol. 89: 980–7

Wittmann T, Gary M, Bokoch GM, Waterman-Storer CM (2003) Regulation of leading edge microtubule and actin dynamics downstream of Rac1. J Cell Biol. 161: 845–851

Yu F, Morin X, Cai Y, Yang X and Chia W (2000) Analysis of partner of inscuteable, a novel player of Drosophila asymmetric divisions, reveals two distinct steps in inscuteable apical localization. Cell 100: 399–409

Yuhui L, Kučera O, Cuvelier D, Rutkowski DM, Deygas M, Rai D, Pavlovič T, Vicente FN, Piel M, Giannone G, Vavylonis D, Akhmanova A, Blanchoin L and Théry M (2022) Compressive forces stabilise microtubules in living cells. BioRxiv doi: 10.1101/2022.02.07.479347

Zarbock A, Deem TL, Burcin TL and Ley K (2007) Galphai2 is required for chemokine-induced neutrophil arrest Blood 110: 3773–3779

Zhang Z, Yu H, Sisi Jiang SJ, Liao J, Lu T, Wang L, Zhang D and Yue W (2015) Evidence for Association of Cell Adhesion Molecules Pathway and NLGN1 Polymorphisms with Schizophrenia in Chinese Han Population. 10.1371/journal.pone.0144719

Zhong M, Clarke S, Vo BT, Khan SA (2012) The essential role of Giα2 in prostate cancer cell migration. Mol Cancer Res. 10: 1380–8.

