## Supplementary figures and images for "Gαi2 Interaction with EB1 Controls Microtubule Dynamics and Rac1 Activity in *Xenopus* Neural Crest Cell Migration"

### Supplementary Figure S1

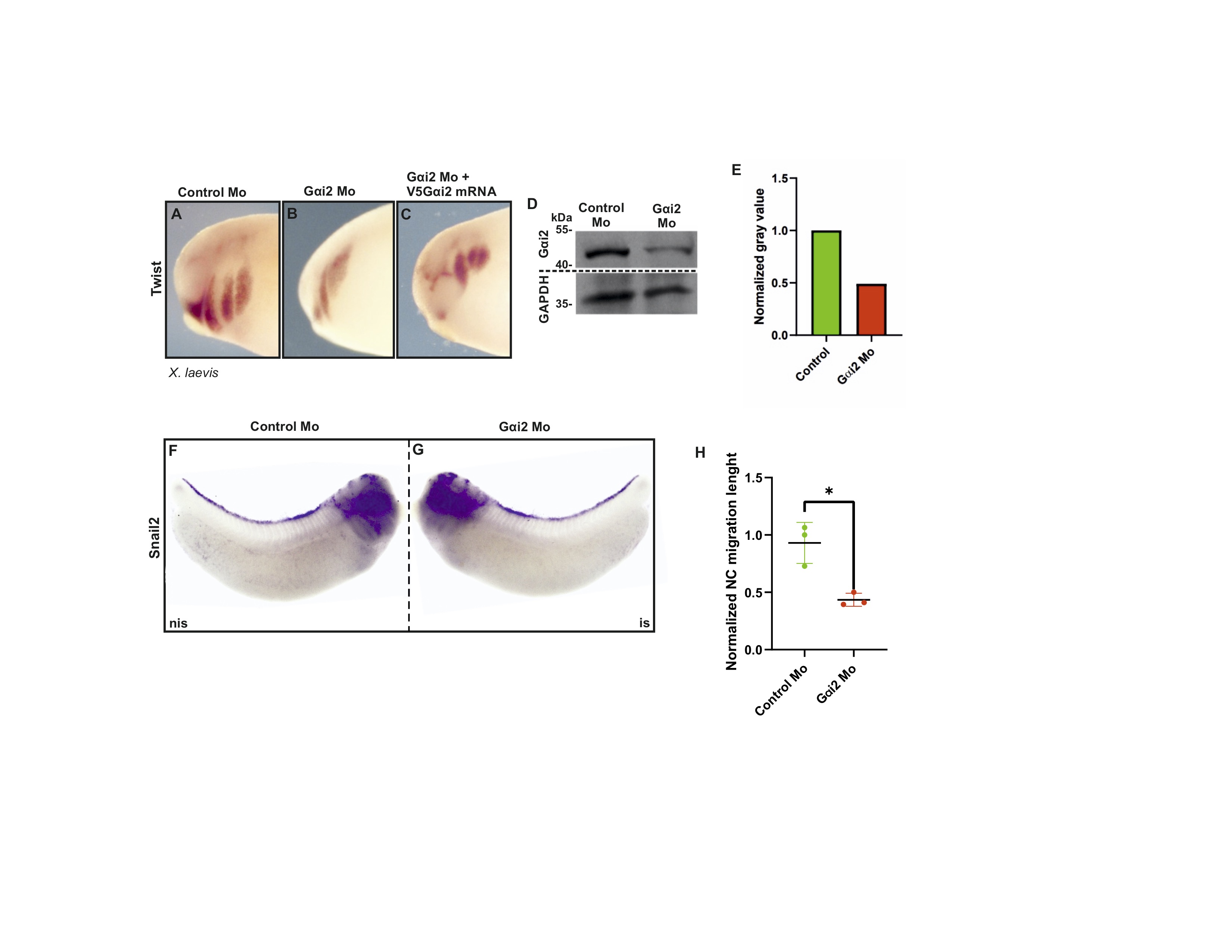

### Supplementary Figure S2

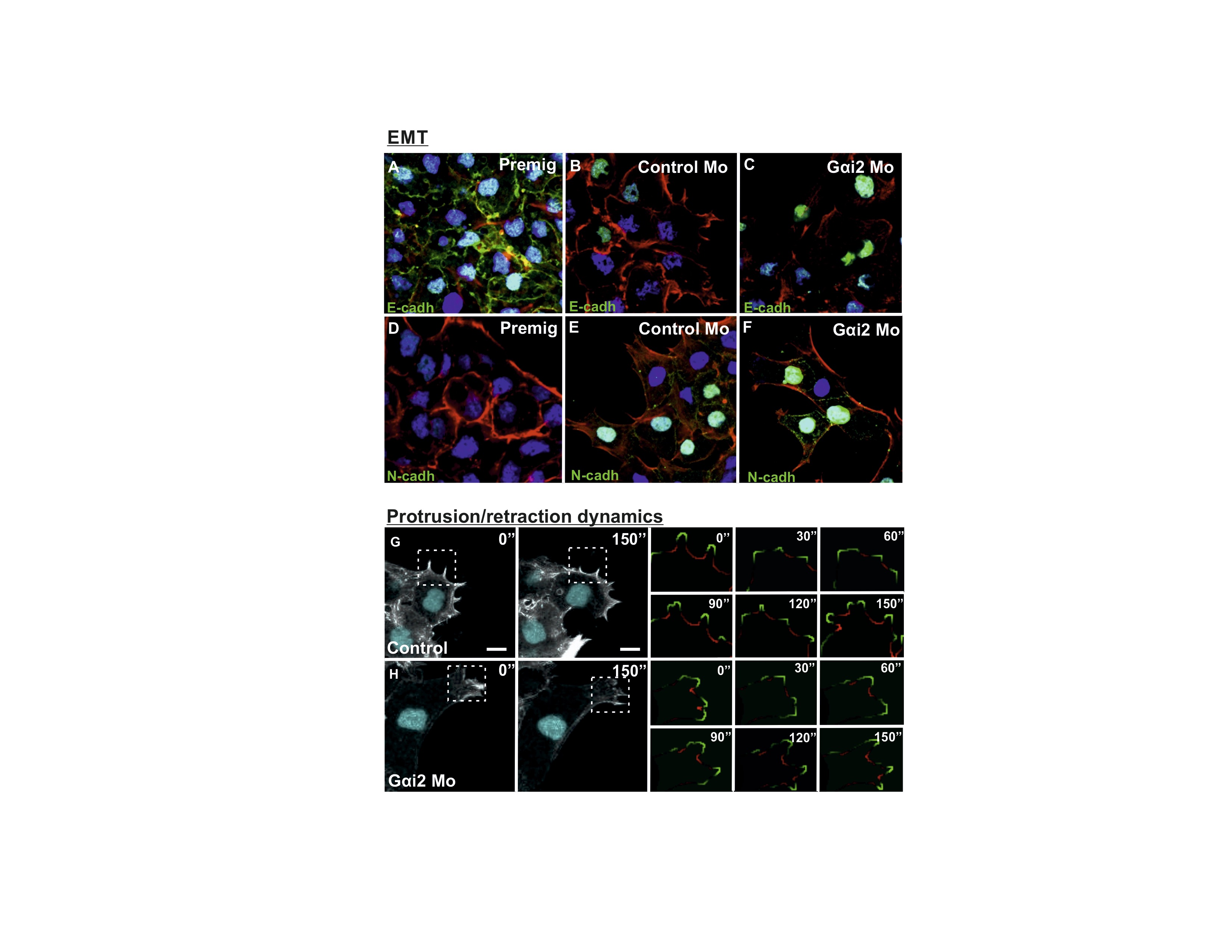

### Supplementary Figure S3

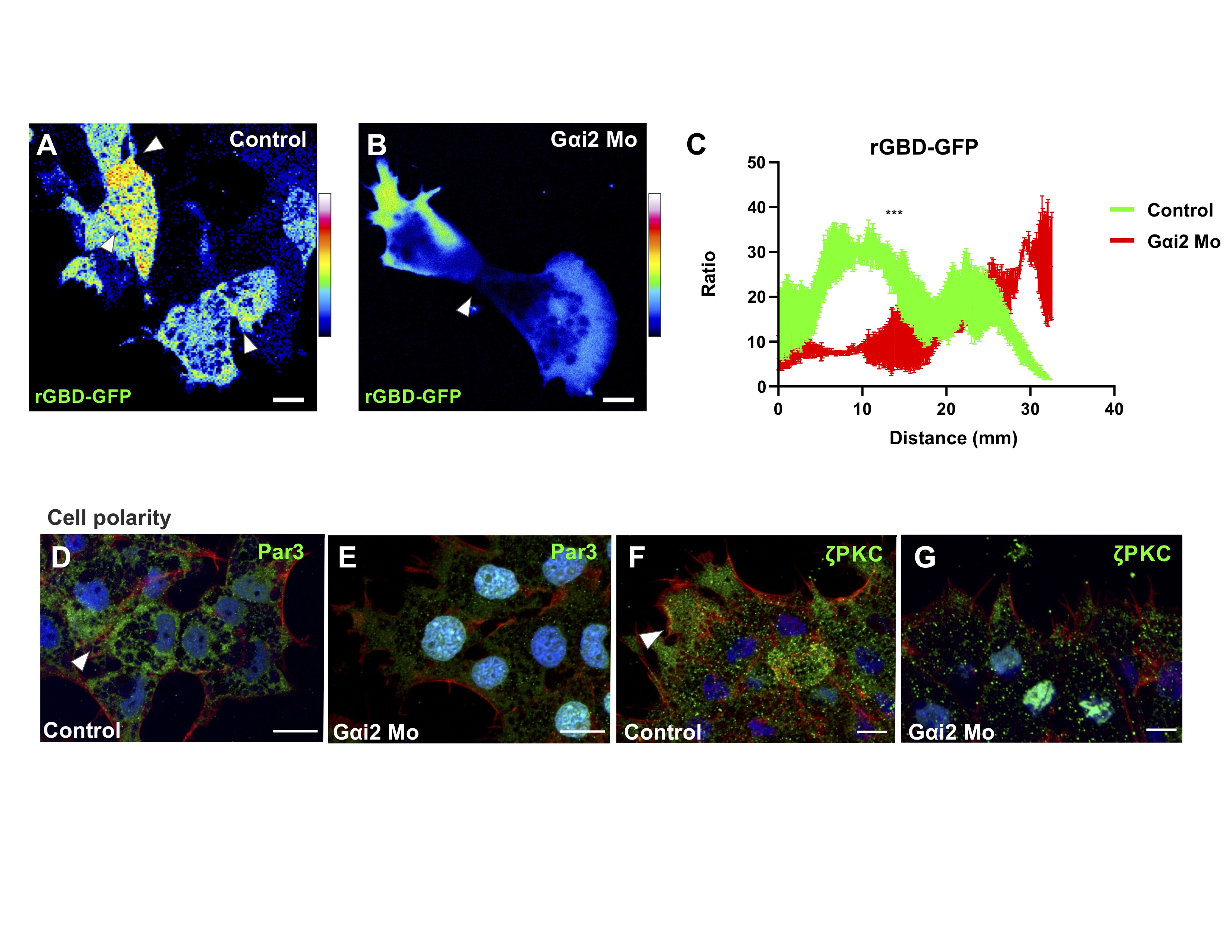

### Supplementary Figure S4

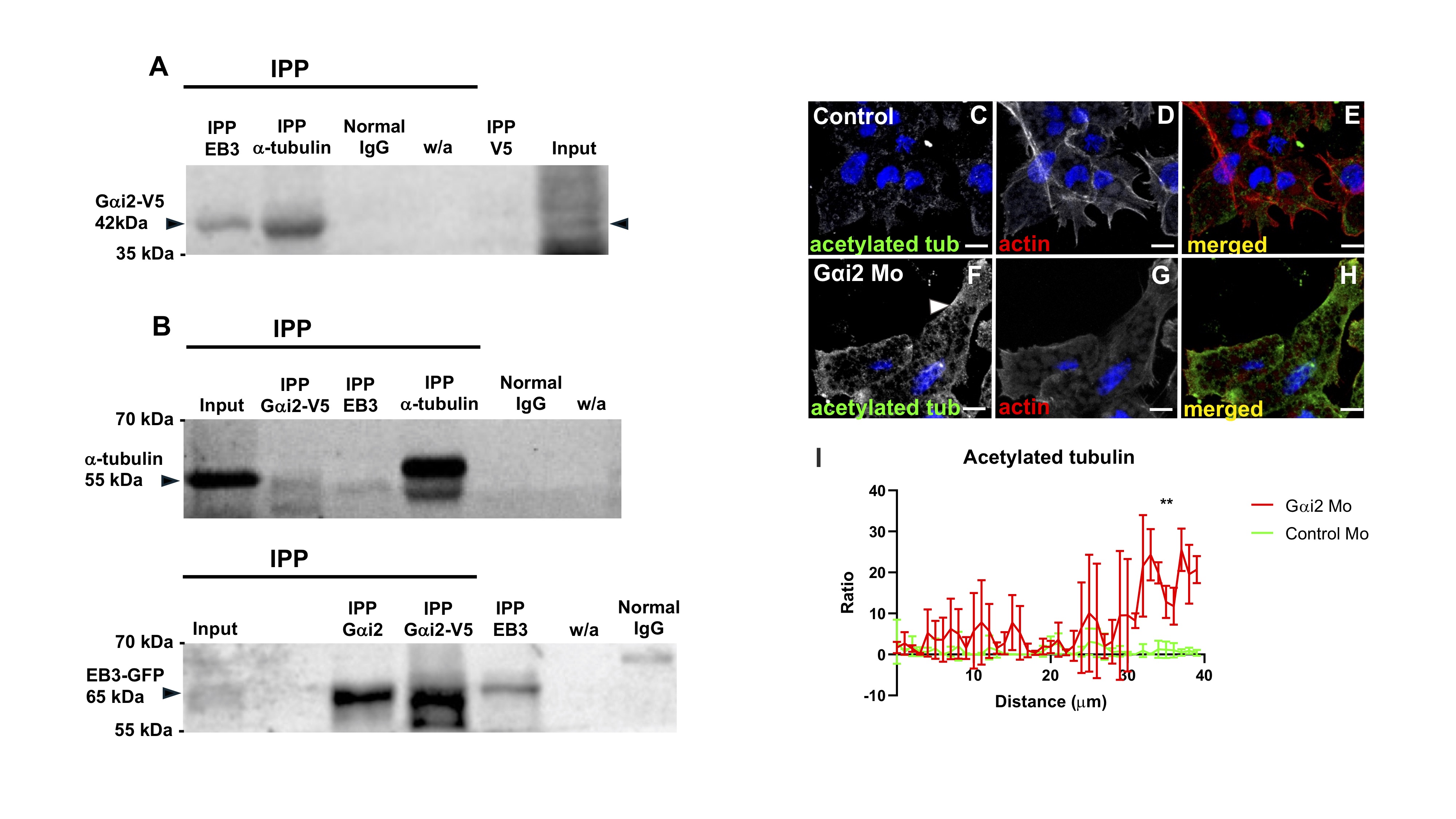
